# Glucagon receptor signaling is indispensable for the healthspan effects of caloric restriction in aging male mice

**DOI:** 10.1101/2025.05.13.653849

**Authors:** Kassandra R. Bruner, Isabella R. Byington, Tyler J. Marx, Anastasiia Vasileva, Temara Fletcher, Susma Ghimire, India J. Zappia, Yashika Shaju, Janan Zeng, Hallie R. Wachsmuth, Thadeus W. Carlyon, David G. Besselsen, Daniel J. Drucker, Frank A. Duca, Jennifer H. Stern

## Abstract

Obesity and type 2 diabetes mellitus accelerate aging, shortening the duration of healthspan. Conversely, chronic calorie restriction (CR) extends healthspan. Research aimed at understanding the mechanism by which CR slows aging has focused heavily on insulin and downstream signaling cascades. Glucagon, a hormone that counter-regulates insulin, is commonly affected by these same interventions. To investigate the role of glucagon in aging we used dietary manipulation, global and liver-specific glucagon receptor knockout, and pharmacological glucagon receptor activation. We found that globally eliminating glucagon receptor signaling (Gcgr KO) decreases median lifespan by 35% in lean mice. These lifespan shortening effects are more robust in diet-induced obese mice (54%). Extending these findings to metabolic health, we found that glucagon receptor signaling is indispensable to the metabolic response to chronic CR in young and aged mice. While CR decreased liver fat, serum triglyceride, and serum cholesterol in WT mice, these metabolic benefits were absent in Gcgr KO mice. In line with these observations, we found that critical nutrient sensing pathways known to improve aging are dysregulated in mice lacking glucagon receptor signaling at the liver (Gcgr^hep-/-^). Liver-specific deletion of the glucagon receptor decreases hepatic AMP Kinase activation in aging mice, regardless of diet. Further, CR decreases hepatic mTOR activity in WT mice, but not in Gcgr^hep-/-^ mice. Together, these findings propose that glucagon signaling plays a critical role in both normal aging and the lifespan and healthspan extension driven by caloric restriction.

## 1. Introduction

The aging field has largely focused on insulin as the primary driver of both the accelerated aging in obesity and the enhanced lifespan associated with calorie restriction (CR). This focus on insulin has dramatically improved our understanding of the biology of aging, establishing that pharmacological and genetic inhibition of insulin signaling pathways can extend healthspan and lifespan in C. elegans [1, 2], drosophila [3, 4] [5], and mice [6, 7]. However, this focus on insulin has largely limited investigation into other hormones and metabolites that are similarly altered by both calorie restriction and obesity.

Glucagon is increased in response to an extended fast [8–11] and glucagon sensitivity is enhanced in response to caloric restriction [9]. Glucagon signaling at the liver activates AMP-activated protein kinase (AMPK) [12], a second messenger that extends healthspan [13, 14] and inhibits mTOR activity [15, 16]. Inhibition of the mTOR pathway promotes longevity and improves healthspan. In fact, both pharmacologic [17, 18] and genetic [19] inhibition of mTOR increases lifespan and decreases age-related disease. Glucagon action at the liver also activates adenylate cyclase, increasing intracellular levels of cyclic AMP (cAMP) [11, 20–22]. Enhancing cAMP signaling extends lifespan in drosophila [23] and improves healthspan in aged mice [24]. Given that glucagon signaling at the liver activates both the cAMP pathway and the IP3-AMPK signaling pathway, this provides two potential signaling pathways [11, 12, 22, 25] by which glucagon receptor signaling may regulate aging.

Several research groups and now the pharmacological industry have shown the potential of glucagon containing di- and tri-agonists to treat obesity and diabetes [26–30]. Pharmacological options that increase glucagon receptor signaling are in development and have established success in clinical trials. In turn, these glucagon receptor agonists are likely to become commercially available soon, encouraging studies to better understand the potential impact of glucagon receptor activity on aging.

Given that glucagon receptor signaling at the liver promotes changes in the same energy sensing pathways (AMPK and mTOR) regulated by CR and glucagon receptor signaling plays a critical role in maintaining both lipid and glucose homeostasis during an extended fast, we aimed to test the hypothesis that glucagon receptor signaling is necessary for both normal healthspan and the healthspan extension resulting from CR.

## 2. Materials and Methods

### 2.1 Animals

All animal procedures in this study were approved by the Institutional Animal Care and Use Committee of the University of Arizona (IACUC protocol 18-478). All experimental procedures were performed according to the National Institutes of Health Guide for the Care and Use of Laboratory Animals. Mice were singly housed and maintained on a 14-hour light/10-hour dark cycle in the University of Arizona Health Sciences temperature (22– 24°C) and humidity (40– 60%) controlled vivarium. Mice were assigned to respective diets at 4 months of age. *Ad libitum*-fed (AL) mice had unrestricted access to chow (NIH-31). Calorie-restricted mice were given either 95% (5% CR), 85% (15% CR), or 60% (40%CR) of *ad libitum* food intake in tablet form (NIH-31, LabDiet; Richmond, Indiana) once daily at 5pm, prior to the onset of the dark cycle. The level of CR was calculated as a function of daily food intake assessed in *ad libitum* mice at 4 months of age (Figure S1A). Diet-induced obese mice were provided *ad libitum* access to high-fat diet, 60% kcal from fat (High Fat Diet; Envigo, TD.06414).

Food intake in AL fed mice was assessed at 4, 6, 8, and 12 months of age to ensure the correct level of restriction was provided (Figure S1). Food intake was assessed in individually housed mice over a 5-day period by weighing hopper feed daily at 9am. We accounted for spillage by weighing any feed at the bottom of each cage.

Global glucagon receptor knockout mice (Gcgr KO), generated as previously described [31], were a gift from Dr. Maureen Charron. To generate Gcgr KO and wildtype littermates (WT), we bred heterozygous (Gcgr^+/-^) males to Gcgr^+/-^ females. Floxed Gcgr mice, generously provided by Dr. Daniel Drucker, were generated as previously described [32]. Albumin-Cre mice were purchased from Jackson Laboratories (Strain #003574). To generate hepatocyte-specific glucagon receptor knockout mice Gcgr^hep-/-^ and wildtype littermate controls, we bred Albumin-Cre^+/-^, Gcgr^Flox/WT^ males to Albumin-Cre^-/-^, Gcgr^Flox/WT^ females. Studies investigating the effects of pharmacologic glucagon receptor agonism on liver cAMP and mTOR activity in at 6- and 17-months of age were performed wildtype male C57BL/6J mice purchased from Jackson laboratories at 4- and 15-months of age (Strain #000664). Mice were allowed to acclimate for 4 weeks prior to initiating glucagon receptor agonism treatment at 5- and 16- months of age.

### 2.2 Lifespan Study

All mice were singly housed and aside from daily visual health checks and bi-monthly body weight monitoring, mice were not disturbed until they reached IACUC-approved criteria for moribundity or were found dead. Because C57BL/6J mice have the propensity to develop dermatitis with age, at the first sign of dermatitis hindlimb nails were clipped weekly and the skin was treated with 2% chlorohexidine ointment daily until the lesions were healed as preventative care. Mice were considered moribund if they could not reach food or water or were unresponsive to external stimuli. Immediately upon discovering an animal that died, a necropsy was performed by a board-certified veterinary pathologist. The carcass was visually examined for external lesions. The abdominal and thoracic cavities were opened with a ventral approach, the oral cavity was opened by cutting through the mandibular rami, and the brain was removed from the skull following removal of the dorsal cranium. All organs were grossly evaluated, and gross lesion descriptions were recorded. Representative tissue samples and all gross lesions were collected into 10% neutral buffered formalin for routine tissue processing and H&E slide preparation, with histopathologic interpretation by the veterinary pathologist.

### 2.3 Body Composition and Indirect Respiration Calorimetry

Body weights were measured after a four hour fast. Prior to initiating calorimetry measurements, fat mass and lean mass was assessed via NMR (EchoMRI™, Houston, TX). Percent of fat mass was calculated based on total body weight. To assess whole body energy expenditure (EE) and substrate utilization (RQ), 6- and 12- month-old mice were single-housed in the Sable Systems International Promethion Core^TM^ 8-Cage metabolic monitoring System (Sable Systems; Las Vegas, NV) within in an environmental chamber maintained at 22°C and 40% humidity on a 12-h light/dark cycle. Mice were first given 48 h to acclimate in the system, followed by an additional 48 hours of data collection. The respiratory quotient (RQ) was calculated as the ratio of volume of CO_2_ produced to the volume of O_2_ consumed Energy expenditure data was calculated using the modified Weir equation: EE(kJ) = (16.5kJ/L X V_O2_) + (4.63kJ/L X V_CO2_) [33]. Data was analyzed using the ExpeData-P and Macro Interpreter software programs (Sable Systems; Las Vegas, NV).

### 2.4 Oral Glucose Tolerance Testing (OGTT) and Oral Glucose Stimulated Insulin Secretion (OGSIS)

Following a 4 h fast starting at 9 AM (Zeitgeber time 4 (ZT 4)), we orally gavaged mice with glucose (2.5 g/kg; 0.1 mL/10 g body weight) (D-Glucose; Fisher Chemical^TM^, Waltham, Massachusetts). All OGTTs began at 1 pm and whole blood glucose concentration was assessed by glucometer (CONTOUR®NEXT glucometer; Bayer Healthcare, Leverkusen, Germany) at 0, 15, 30, 60, 90, and 120 minutes after glucose gavage. Blood for serum insulin (oral glucose stimulated) and glucose determination was collected from the tail vein at baseline prior to glucose gavage and 15 minutes after glucose administration [34, 35].

### 2.5 Insulin Tolerance Testing

To assess differences in insulin sensitivity, 4h fasted mice were injected intraperitoneally with insulin (0.25 IU/kg body weight). This lower concentration of insulin was used to prevent hypoglycemia in calorie restricted mice with decreased basal glucose and improved insulin sensitivity [34]. Blood glucose concentration was assessed using glucometer at baseline and 15, 30, 60, 90, and 120 minutes after insulin injection.

### 2.6 Pharmacologic Treatment with a long-acting glucagon analogue (GCGA)

The long-acting glucagon analogue NNC9204-0043 was provided to us by the Novo Nordisk Compound Sharing Program (Novo Nordisk, Denmark).

#### Acute GCGA treatment (single dose)

At 9 AM (Zeitgeber time 4 (ZT 4)), 6-month-old male wildtype C57BL/6J mice (Jackson Laboratories, Strain #000664) were given a single injection of either PBS or a long-acting glucagon analogue (GCGA; 1.5 nmol/kg BW subcutaneous, NNC9204-0043 Novo Nordisk, terminal half-life 5-6 hours). Food was removed from the cage at time of injection and 4 hours later, livers and serum were collected.

#### Chronic GCGA treatment (4 weeks of three times weekly injections)

At 5 and 16 months of age, male wildtype C57BL/6J mice (Jackson Laboratories, Strain #000664) were assigned to either treatment with a long-acting glucagon analogue (GCGA; 3 nmol/kg BW subcutaneous, NNC9204-0043 Novo Nordisk, terminal half-life 5-6 hours) or PBS vehicle control for four weeks (three times per week). At completion of the one-month treatment period and 24 hours after the last GCGA dose, mice were fasted for 4 hours and livers were collected.

### 2.7 Tissue Collection

4h fasted mice were sacrificed at Zeitgeber time 8 by decapitation after acute exposure to high dose isoflurane. Trunk blood was collected into a 1.7 mL microcentrifuge tube and allowed to clot on ice for 30 minutes. Serum was collected and stored at -80°C in 50µL aliquots after centrifugation at 3,000 x g for 30 minutes. Tissues were snap frozen on dry ice and stored at -80°C until analysis. Frozen livers were powdered with a liquid nitrogen cooled mortar and pestle then stored at -80°C in preparation for liver tissue analyses.

### 2.8 Hepatic triglyceride content

15-20mg of frozen powdered liver tissue was weighed and recorded prior to sonicating in 100uL ice cold PBS. Next, 1mL of 100% ethanol was added to each sample to extract lipid. Samples were then vortexed vigorously for 20 minutes and centrifuged at 1,600 x g at 4°C. Supernatant was transferred to a fresh tube for final analysis of liver triglycerides (Cat. # T7531, Pointe Scientific Inc., Canton, MI). Liver triglyceride content was calculated based on grams of liver assayed [34, 35].

### 2.9 Serum hormone and metabolite assays

We analyzed serum glucose, triglyceride, and total cholesterol concentrations using enzymatic colorimetric assays (Glucose: Cat. # G7519, Pointe Scientific Inc., Canton MI; triglyceride: Cat. # T7531, Pointe Scientific Inc., Canton, MI; and total cholesterol: Cat # TR13421, Thermo Scientific™, Middletown, VA). We quantified serum insulin using a commercially available enzyme-linked immunosorbent assay (Cat. # 80-INSMSU-E10, Alpco, Salem, NH). We calculated the Homeostatic Model Assessment for Insulin Resistance (HOMA-IR) index using the formula: HOMA-IR=fasting glucose in mmol/l*fasting insulin in μU/ml/22.5 [36].

### 2.10 Liver Glycogen and cAMP Content

We quantified liver glycogen content using the colorimetric assay as described by Lo and colleagues (1970) [37]. 10-15 mg of powdered liver frozen at -80°C was weighed before being boiled and shaken in 30% KOH saturated with NA_2_SO_4_ for 30 minutes. We then precipitated glycogen with 95% ethanol and pelleted by centrifugation at 3000 x g for 30 minutes. The supernatants were aspirated, and glycogen pellets dissolved in distilled H_2_O. 5% Phenol was added to the samples followed by a rapid, forceful stream of H_2_SO_4_. Samples were then incubated at 30°C for 20 minutes and absorbance was read at 490 nm. Glycogen content was expressed per gram of liver tissue initially weighed.

Liver cAMP was quantified using the Enzo Direct cAMP enzyme-linked immunosorbent assay (ELISA), Cat #ADI-901-066A (Enzo Life Sciences, Farmingdale, NY). Prior to assay, frozen powdered liver was weighed and quickly sonicated in 10 volumes of ice cold 0.1M HCl. The homogenate was centrifuged at 600 x g at 4°C for 10 minutes. Supernatant was removed and immediately assayed by ELISA. Liver cAMP content was expressed per gram of liver tissue initially weighed.

### 2.11 Western Blotting

To assess mTOR and AMPK activation, we quantified the expression of mTOR (phosphorylated and total) and AMPK (phosphorylated and total) proteins in liver lysates. Briefly, protein lysates were prepared by sonicating frozen powdered liver tissue in ice cold RIPA buffer (Cat# sc-24948A; Santa Cruz Biotechnology, Dallas, TX) combined with Halt protease and phosphatase inhibitor cocktail (Cat # 78442 Thermo Scientific, Waltham, MA). Samples were centrifuged at 4°C, supernatants removed, aliquoted, and quickly stored at -80 for no longer than 3 weeks prior to western blotting. Total protein was quantified using the Pierce™ Rapid Gold BCA Protein Assay (Cat # A55860 Thermo Scientific, Waltham, MA). 40 µg protein per sample was loaded and separated on a 4-12% bis tris gel (Cat #NP0321BOX, Invitrogen/ Thermo Scientific, Waltham, MA) and transferred onto a nitrocellulose membrane (0.2 µm pore size) using the Trans-Blot Turbo Transfer System (Bio-Rad; Hercules, CA). Membranes were blocked for 1h at room temperature in a blocking solution [5% w/v nonfat dry milk in Tris-buffered saline with 0.5% Tween 20 (TBST)], then incubated overnight at 4°C with primary antibodies diluted to 1:1000 in blocking solution. The following day, membranes were washed five times for 5 minutes with TBST, then incubated with a secondary antibody (Anti-rabbit IgG, HRP-linked Antibody, Cell signaling #7074) diluted 1:2000 in blocking solution for 1h at room temperature. Signals were visualized using Thermo Scientific™ SuperSignal™ West Pico PLUS Chemiluminescent Substrate (Thermo Cat # PI34578) and imaged in the Azure 600 Western Blot Imaging System (Azure; Dublin, CA). All images were obtained using the same exposure time within each protein and band intensities quantified using the AzureSpot Pro image analysis software (Azure; Dublin, CA). All antibodies were purchased from Cell Signaling (AMPK Rabbit mAb: Cat #5831S, Phospho-AMPK (Thr172) Rabbit mAb: Cat #2535S, S6 Ribosomal Protein Rabbit mAb: Cat # 2217S, Phospho-S6 Ribosomal Protein (Ser240/244) XP® Rabbit mAb: Cat # 5364S (Cell Signaling, Danvers, MA).

### 2.12 RNA Extraction and mRNA expression of hepatic genes

We extracted RNA from frozen powdered livers using TRIzol™ Reagent. (Thermo Fisher Scientific, Waltham, MA) following manufacturer protocol. Immediately following extraction, RNA was washed with water-saturated butanol and ether (3 times each) using the method of Krebs et al. (2009) to eliminate phenol contamination [38]. We performed reverse transcription with iScript cDNA synthesis kit (Bio-Rad Laboratories, Hercules, CA) and RT-qPCR using SsoAdvanced Universal SYBR® Green Supermix (Bio-Rad Laboratories, Hercules, CA) on the Applied Biosystems QuantStudio 3 Real-Time PCR System (Applied Biosystems™, Foster City, CA). We analyzed raw Ct values using LinReg PCR analysis software to determine amplification efficiency [39]. Genes of interest were normalized to mouse Tbp (TATA-box binding protein) mRNA expression and the fold change in gene expression was calculated using the efficiencyΔΔCt method [40]. Fold change for all genotype and diet groups was calculated against the ad libitum-fed WT group within each age group (6 or 17 months of age). Mouse primer sequences for all genes are presented in Table 1.

**Table 1.**
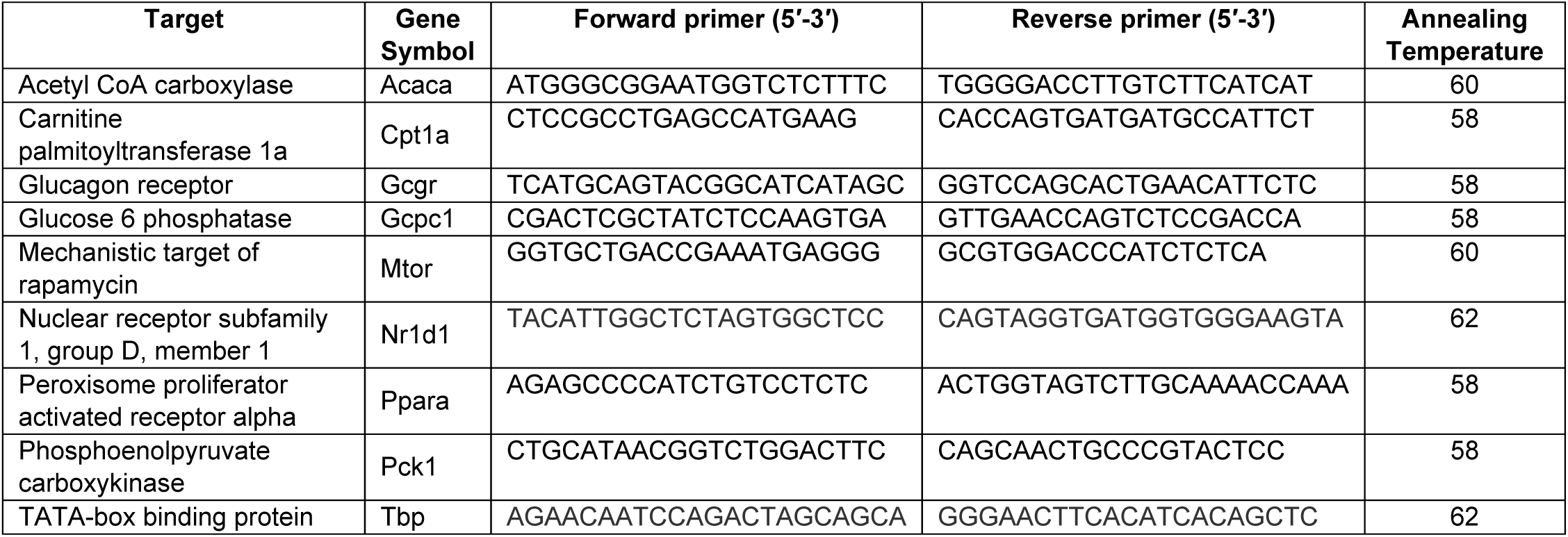
List of Mus musculus Primer Sequences for RT-PCR.

### 2.13 Assessment of Physical Function

To assess balance and coordination as an indicator of physical function, we utilized a continuous-acceleration apparatus (Rota-rod; Rotamex-5 Columbus Instruments, Columbus, OH) on which mice were placed while stationary. Once mice stabilized their posture, incremental rod acceleration was initiated starting at 4 rotations per minute (rpm) and increasing by 0.5 rpm every 5 seconds. We assessed time to fall (seconds) onto a foam cushion underneath the apparatus. Three trials per mouse were completed and each data point represents the average time to fall per mouse.

### 2.14 Statistics

We performed statistical analyses in GraphPad Prism Version 10.4.2 (GraphPad Software, San Diego, California, USA). We assessed survival curves using the Gehan-Breslow-Wilcoxon test because of its sensitivity to early mortality differences [41]. For all cross-sectional studies in 6- and 17-month-old Gcgr KO versus WT the interaction between the two on all dependent variables. The probability of difference between means was assessed after a Tukey’s adjustment for multiple comparisons. For studies examining the effects of GCGA in male wildtype C57BL/6J mice, independent t-tests were performed within each age (6- or 17-months of age) to determine the effects of glucagon receptor agonism on all outcome variables. Raw data were plotted in GraphPad Prism Version 10.4.2 for Windows (GraphPad software). All data are presented as mean ± SEM.

## 3. Results

### 3.1 Global deletion of the glucagon receptor decreases lifespan across nutritional paradigms (chow-fed lean, diet-induced obesity, and calorie restriction)

To examine the role of glucagon receptor signaling in normal lifespan and obesity-induced lifespan curtailment, we performed a lifespan study in low-fat chow fed lean, and diet induced obese Gcgr KO and WT littermate male mice. Whole body glucagon receptor deletion decreased median lifespan in ad libitum chow fed mice by 35% (524 days KO vs. 817 days WT, P=0.0099, Gehan-Breslow-Wilcoxon test: Figure 1A). Obesity shortens lifespan more robustly in glucagon receptor knockout than in wildtype mice (Median lifespan of 374 days KO vs. 690 days WT, Gehan-Breslow-Wilcoxon test: P=0.05; **Figure 1A**). Post-mortem histopathological analysis of a subset of low-fat chow fed Gcgr KO mice randomly selected for necropsy revealed several instances of islet cell carcinoma (Supplemental Data Table 1).

**Figure 1:**
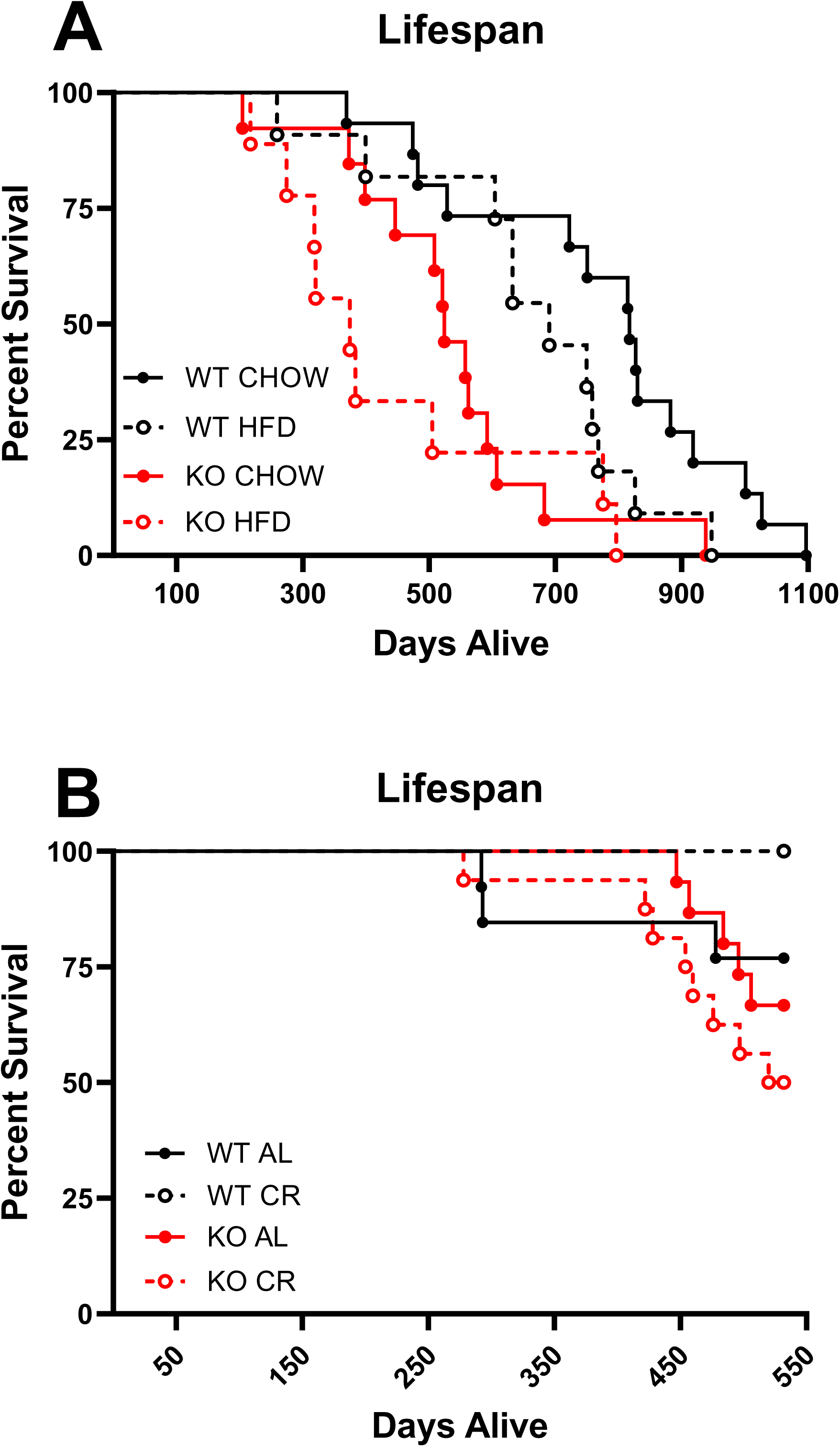
Global deletion of glucagon receptor decreases lifespan. Survival curves of A) Ad libitum chow-fed and high fat diet (HFD) fed mice lacking glucagon receptors (Gcgr KO) have a shorter lifespan compared to wild type littermates (WT) (CHOW Median Lifespan: 524 days KO vs. 817 days WT, P=0.0099, Gehan-Breslow-Wilcoxon test; HFD Median lifespan: 374 days KO vs. 690 days WT, P=0.05 Gehan-Breslow-Wilcoxon test). Chow: n= 15 WT and 13 KO, HFD n= 11 WT and 9 KO mice). Gcgr KO: global glucagon receptor knockout, WT: wildtype littermate controls. B) Decreased survival up to 19 months of age in calorie-restricted (CR; 15% since 4 Mo.) Gcgr KO mice compared to WT littermates (P = 0.008, Gehan-Breslow-Wilcoxon test). Ad libitum (AL), 15% calorie restricted (15% CR).

Because caloric restriction increases lifespan, we monitored lifespan in a subset of our aging mice maintained on CR restriction through 19 months of age. At 19 months of age, only 50% of calorie restricted (15% restriction) GKO mice remained alive, while 100% of calorie restricted wildtype mice were alive (P= 0.008, Gehan-Breslow-Wilcoxon test; **Figure 1B**).

### 3.2 Body weight and fat mass is decreased in aged Gcgr KO mice, despite no effect on food intake and minimal changes in whole body energy metabolism

Despite the shortening of lifespan in mice lacking glucagon receptor signaling, chow-fed GKO mice resist body weight (**Figure 2A and D**) and fat mass (**Figure 2E**) gain during middle age (12 months). In fact, ad libitum (AL) fed Gcgr KO mice maintain lower fat mass compared to WT littermates at 12 months (P<0.001, **Figure 2E**), not affect body weight, but did decrease fat mass (P=0.0001, **Figure 2C**). Despite maintaining a lower body weight and fat mass, global deletion of Gcgr did not affect ad libitum food intake at 4, 6, or 12 months of age (**Supplemental Figure 1A-C**). Further, global deletion of Gcgr did not affect whole body energy expenditure adjusted for lean mass (EE, **Supplemental Figure 2A-F**), or substrate utilization (RQ, **Supplemental Figure 2G-L**) in AL fed mice at 6 or 12 months of age.

**Figure 2:**
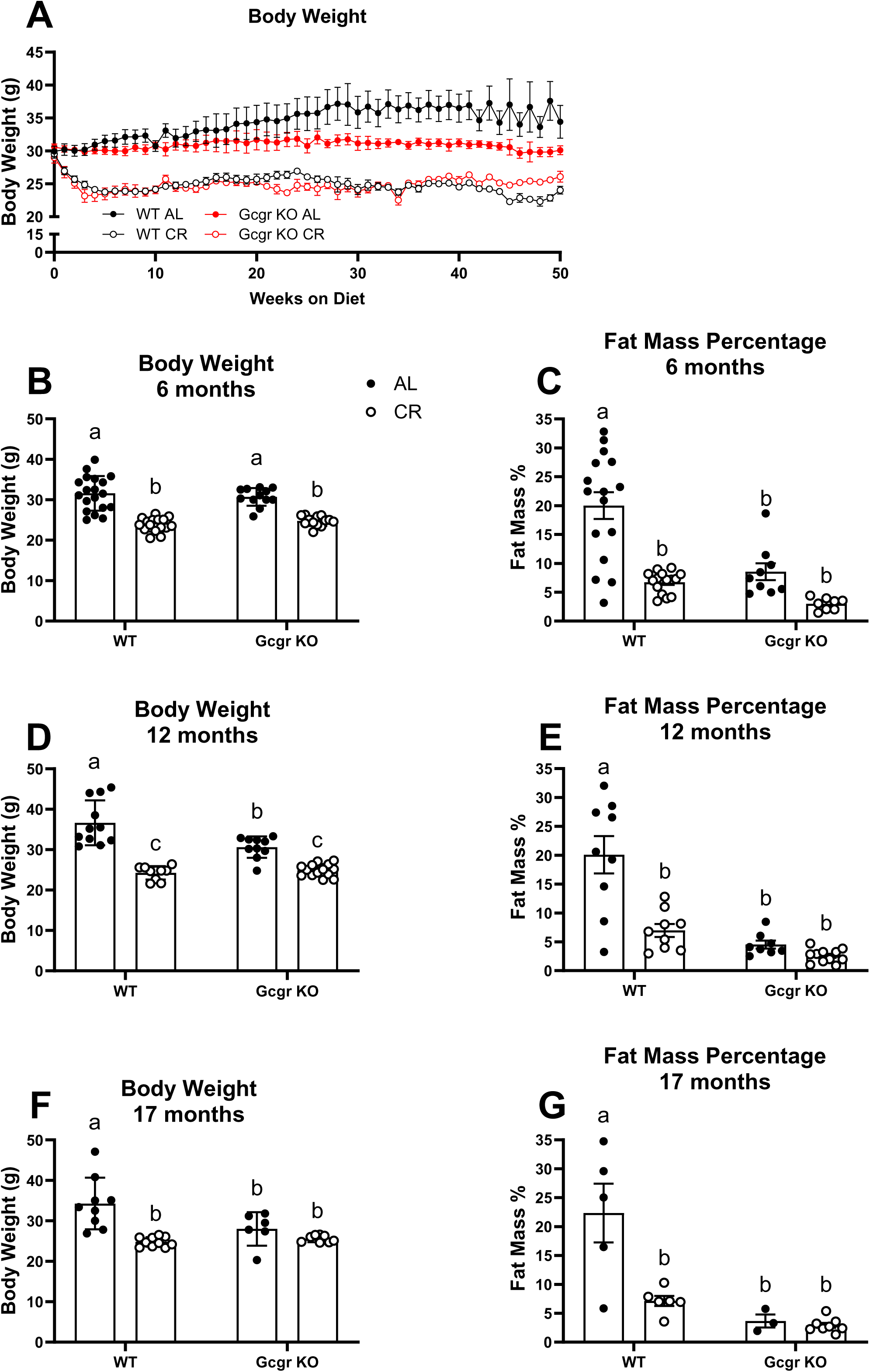
Body weight and fat mass throughout calorie restriction studies. Weekly body weight over 12.5 months of CR initiated at 4.5 months of age. Body weight and fat mass (%) at 6 months of age (B-C; n=12-17 mice per group), 12 months of age (D-E; n=6-10 mice per group), and 17 months of age (F-G; n=5-9 mice per group). Ad libitum (AL), 15% calorie restricted (15% CR). Gcgr KO: global glucagon receptor knockout, WT: wildtype littermate controls. ^a,b^Superscript letters that differ indicate differences, P≤ 0.01; two-way ANOVA with Tukey’s adjustment for multiple comparisons, data are means ± SEM.

We observed a genotype by diet interaction when examining the effects of caloric restriction (15% initiated at 4 months of age, CR) on body weight and fat mass. CR decreased body weight and fat mass in WT 6-, 12-, and 17-month-old mice (P<0.01 at all ages; **Figures 2B-G**). In contrast, GKO mice, with already lowered body and fat mass, only respond to CR with a decrease in body weight at 6 and 12 months of age (P<0.01), with no effect on body weight at 17 months of age and no change in fat mass at 6, 12, or 17 months of age. The whole-body energetic response to caloric restriction was similar between WT and Gcgr KO mice. CR decreased light cycle EE at 12 but not 6 months of age in both WT and Gcgr KO mice (**Supplemental Figure 2B and E**). CR decreased EE and RQ during the dark cycle only at 6 months of age in WT mice (**Supplemental Figure 2C and I**). Indicative of an increase in whole body lipid oxidation during fasting and as we have previously shown [42], CR decreased light cycle RQ at 6 and 12 months of age, regardless of genotype (**Supplemental Figure 2H and K**).

### 3.3 Glucagon receptor signaling is indispensable to the metabolic response to chronic caloric restriction at young adulthood and advanced age in male mice

Given our observation that global glucagon receptor deletion shortens lifespan across nutritional paradigms, we performed a cross-sectional study at 6 and 17 months of age to assess the metabolic response to chronic 15% CR (initiated at 4 months of age) in GKO and WT littermate control mice. We first assessed the impact of global glucagon receptor deletion, CR, and the interaction between genotype and CR on oral glucose clearance and insulin sensitivity. Others have previously demonstrated that global glucagon receptor deletion improves glucose homeostasis and lowers circulating glucose and insulin in ad libitum fed mice [31, 32]. In line with these findings, we found that basal (4h fasted) glucose (P<0.0001, **Figures 3A&E**) and insulin (P<0.01, **Figures 3B&F**), along with HOMA-IR (P<0.001, **Figures 3I&M**) is lowered in AL fed Gcgr KO mice compared to WT littermates at both 6 and 17 months of age. Gcgr deletion decreased oral glucose stimulated insulin at 17-months of age (P<0.05), but not 6-months of age in AL fed mice (**Figures 3I&L**). Because we observed a robust genotype effect on basal glucose, we corrected for basal glucose (Delta OGTT) when assessing glucose clearance (Raw OGTT data provided in **Figures 3C-D and 3G-H**). In AL fed mice, Gcgr knockout improved oral glucose clearance (Delta OGTT) at 6 months of age (P=0.0002), but not 17 months of age (**Figures 3L&P**). Similar to the findings of Gelling and colleagues [31], when corrected for basal glucose levels (Raw ITT data provided in **Supplemental Figures 3A-B and E-F**), glucagon receptor deletion had no effect on insulin sensitivity, as assessed by an insulin tolerance test, regardless of age (**Supplemental Figures 3C-D and G-H**).

**Figure 3:**
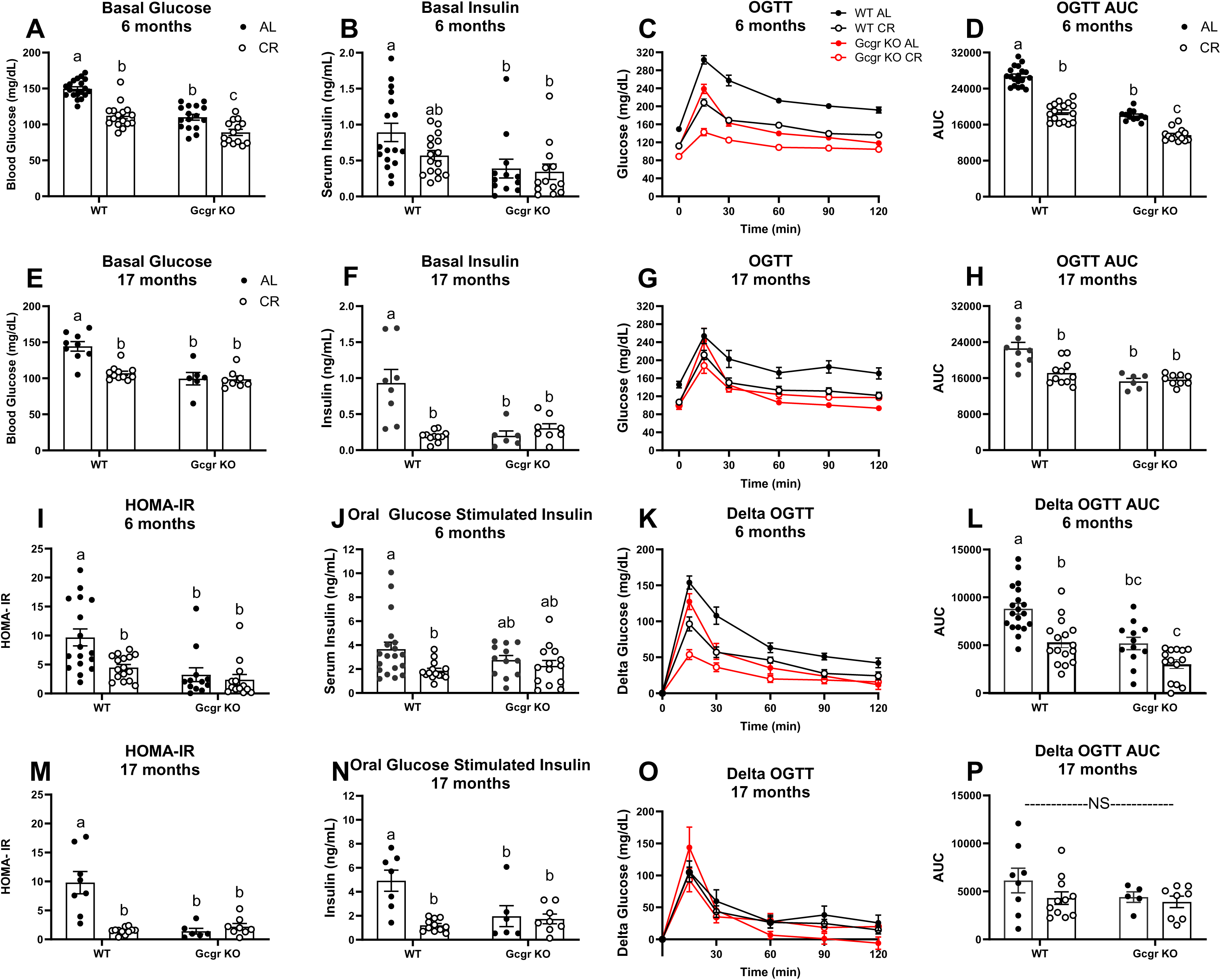
Glucose homeostasis. Basal glucose and insulin at 6 (A-B; n=12-19 mice per group) and 17 (E-F; n=6-11 mice per group) months of age. Oral glucose clearance (2.5g/ kg glucose) and corresponding area under the curve at 6 (C-D) and 17 (G-H) months of age. HOMA-IR (Homeostatic Model Assessment for Insulin Resistance) at 6 (I) and 17 (M) months of age. Oral glucose-stimulated insulin at 6 (J) and 17 (N) months of age. Oral glucose clearance as expressed as change from baseline (Delta OGTT) and corresponding area under the curve at 6 (K-L) and 17 (O-P) months of age. AL, ad libitum fed; CR, calorie restricted (15% initiated at 4.5 months of age). Gcgr KO: global glucagon receptor knockout, WT: wildtype littermate controls. ^a,b^Superscript letters that differ indicate differences, P≤ 0.01; two-way ANOVA with Tukey’s adjustment for multiple comparisons. NS, not significant, data are means ± SEM.

#### Caloric restriction fails to further improve glucose homeostasis in Gcgr KO mice

We previously showed that lifelong 15% CR improves glucose clearance, lowers circulating insulin, and improves insulin sensitivity in aging male mice [34]. In line with these findings, in WT littermates CR decreased basal glucose (P<0.0001, **Figures 3A&E**), HOMA-IR (P<0.01, **Figures 3I&M**), and oral glucose stimulated insulin (P<0.05, **Figures 3J&N**) at both 6- and 17-months of age. In WT mice only, CR robustly decreased basal insulin at 17 months of age (P<0.0001, **Figure 3F**), with a modest, non-significant reduction at 6 months of age (P=0.13, **Figure 3B**). Apart from a decrease in basal glucose at 6 months of age (P=0.002, **Figure 3A**), CR failed to lower basal glucose (**Figure 3E**), insulin (**Figure 3B&F**), oral glucose stimulated insulin (**Figure 3J&N**) and failed to improve oral glucose clearance (**Figures 3L&P**) in mice lacking Gcgr signaling. When corrected for basal glucose levels, neither CR nor genotype affected insulin sensitivity, as assessed by an insulin tolerance test, regardless of age (**Supplemental Figure 3**).

Because liver glycogen stores contribute to glucose homeostasis as the primary source of stored glucose, we examined the impact of both Gcgr deletion and CR on hepatic glycogen stores. Global Gcgr deletion had only minimal impact on liver glycogen stores, with an increase in response to CR at 6 months of age. Neither genotype nor diet affected liver glycogen at 17 months of age (**Supplemental figure 4A-B**).

Although there is some debate as to whether glucagon’s glucose-mobilizing effect during a fast is primarily mediated though cAMP [11, 43], it is well-established that glucagon action at the liver increases cAMP production [11, 20–22]. Thus, we next quantified hepatic cAMP concentrations and found that neither genotype nor CR affected liver cAMP content at 6 or 17 months of age. (**Supplemental figure 4C-D**). Because we suspected that glucagon stimulated changes in liver cAMP production may only be detectable after acute receptor activation, we next tested the acute and chronic effects of exogenous glucagon receptor agonism on liver cAMP content in wildtype male C57BL/6J mice. While a single dose of a long-acting glucagon analogue (GCGA; 1.5 nmol/kg BW subcutaneous, NNC9204-0043 Novo Nordisk) resulted in a rise in liver cAMP 4 hours post injection (P=0.005, **Supplemental Figure 5A**), there was no effect in mice sacrificed 24h after their last dose of daily, one month GCGA treatment (three times per week; 3nmol/kg BW subcutaneous, NNC9204-0043 Novo Nordisk) at 6 or 17 months of age (**Supplemental Figure 5B-C**).

#### Caloric restriction fails to improve lipid homeostasis in Gcgr KO mice

Our previous work demonstrated that lifelong 15% CR decreases hepatic lipid accumulation in 18-month-old aging mice [34]. Because glucagon signaling at the liver regulates lipid homeostasis by stimulating fatty acid oxidation and inhibiting de novo lipogenesis [8, 26], we hypothesized that GKO mice would be resistant to the lipid lowering effects of CR. In line with our previous findings, CR decreased liver triglyceride content, serum triglyceride, and serum cholesterol at 6 (P≤0.01, **Figures 4A-C**) and 17 months of age (P≤0.05, **Figures 4D-F**) in WT littermates. Mice that globally lack glucagon receptor fail to respond to chronic calorie restriction with a decrease in liver and serum triglyceride and serum cholesterol at both 6- (**Figures 4A-C**) and 17- (**Figures 4D-F**) months of age. Interestingly, at 17 months of age, liver triglyceride content (P=0.006, **Figure 4D**) and circulating triglyceride (P=0.036, **Figure 4E**) are decreased in AL fed GKO mice compared to AL fed WT littermates. CR fails to further decrease liver and serum triglyceride in 17-month-old GKO mice (**Figures 4D-E**).

**Figure 4:**
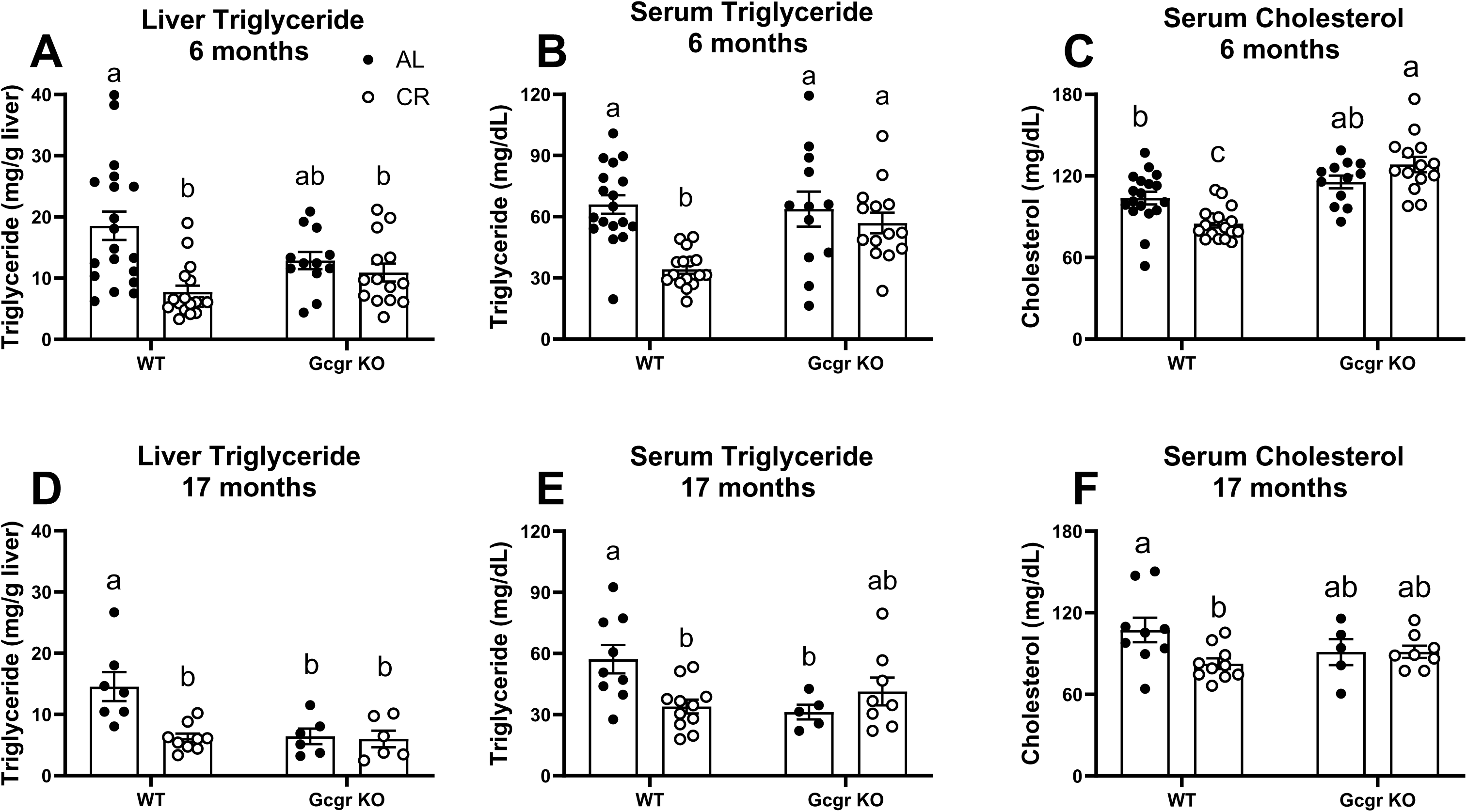
Lipid homeostasis. Liver triglyceride content, serum triglyceride, and serum cholesterol at 6 months of age (A-C: n= 12-19 mice per group) and 17 months of age (D-F: n= 5-11 mice per group) AL, ad libitum fed; wildtype littermate controls. ^a,b^Superscript letters that differ indicate differences, P≤ 0.01 (6 month old) and P≤0.05 (17 month old); two-way ANOVA with Tukey’s adjustment for multiple comparisons, data are means ± SEM.

To investigate potential differences in the mRNA expression of genes that encode for enzymes and signaling molecules that regulate glucose and lipid metabolism at the liver, we performed qPCR on livers collected from AL and CR fed Gcgr KO and WT mice. We first confirmed barely detectable mRNA expression of the glucagon receptor in both 6- and 17-month-old Gcgr KO mice compared to WT littermates, regardless of diet (P<0.0001, **Supplemental Figures 6A&E**). We found little to no effect of genotype or diet on the mRNA expression of most genes, except for the gluconeogenic genes, Phosphoenolpyruvate Carboxykinase (Pck1) and Glucose 6 Phosphatase (G6pc1), and the circadian clock gene Nr1d1 (also known and REV-ERBα). Glucagon receptor deletion decreased Pck1 mRNA expression in 6-month-old calorie restricted mice compared to WT (P<0.05, **Supplemental Figure 6B**). G6pc1 mRNA expression decreased in response to CR, independent of genotype, but only at 6 months of age (P<0.05, **Supplemental Figures 6C&G**). The expression of Nr1d1, a circadian clock gene that can regulate both lipid and glucose metabolism (refs) decreased in response to calorie restriction at both 6- (P=0.028) and 17- (P=0.023) months of age in WT littermates, with no effect in Gcgr KO mice (**Supplemental Figures 6L&P**).

### 3.4 Nutrient sensing pathways that regulate aging are dysregulated in mice lacking glucagon receptor signaling at the liver

The liver is the main site of glucagon action [8, 35, 44]. Therefore, we next set out to assess the effects of hepatic glucagon receptor signaling, caloric restriction, and the interaction between genotype and diet on two nutrient signaling pathways at the liver known to regulate aging; mechanistic target of rapamycin (mTOR) and AMP-activated protein kinase (AMPK) [13, 14, 17–19, 45]. While increasing mTOR signaling accelerates aging [46], inhibition of mTOR activity slows aging [17, 18]. Increasing AMPK activation can also slow aging [13, 14]. We studied the response to chronic caloric restriction (initiated at 4 months of age) in male 17-month-old hepatocyte-specific glucagon receptor knockout mice (Gcgr^Hep-/-^) and wildtype littermates (WT) under three dietary conditions: AL fed, 5% CR, or 40% CR. We assessed mTORC1 activity by quantifying the ratio of phosphorylated (Ser^444/440^) to total ribosomal S6 protein, a common surrogate for mTORC1 activity [46, 47]. Within WT littermates, 5% CR was not sufficient to decrease mTOR activation. However, a 40% restriction suppressed mTOR activation, as assessed by the phosphorylation of S6 (P=0.02, **Figure 5A-B**). In contrast, Gcgr^Hep-/-^ mice fail to respond to CR with a decrease in mTOR activity, suggesting that glucagon signaling at the liver is required for this nutrient sensing pathway to respond to chronic caloric restriction. While there was no effect of diet on AMPK activation, we observed a robust genotype effect on AMPK activation (P=0.0002). Regardless of diet, liver-specific deletion of glucagon receptor decreased the activation of AMPK, as assessed by the phosphorylation of threonine^172^ (**Figures 5C-D**, full blots provided in Supplemental Figure 8A and 8B). Together, these data suggest that glucagon receptor signaling specifically at the liver regulates nutrient sensing pathways that affect healthspan and aging.

**Figure 5:**
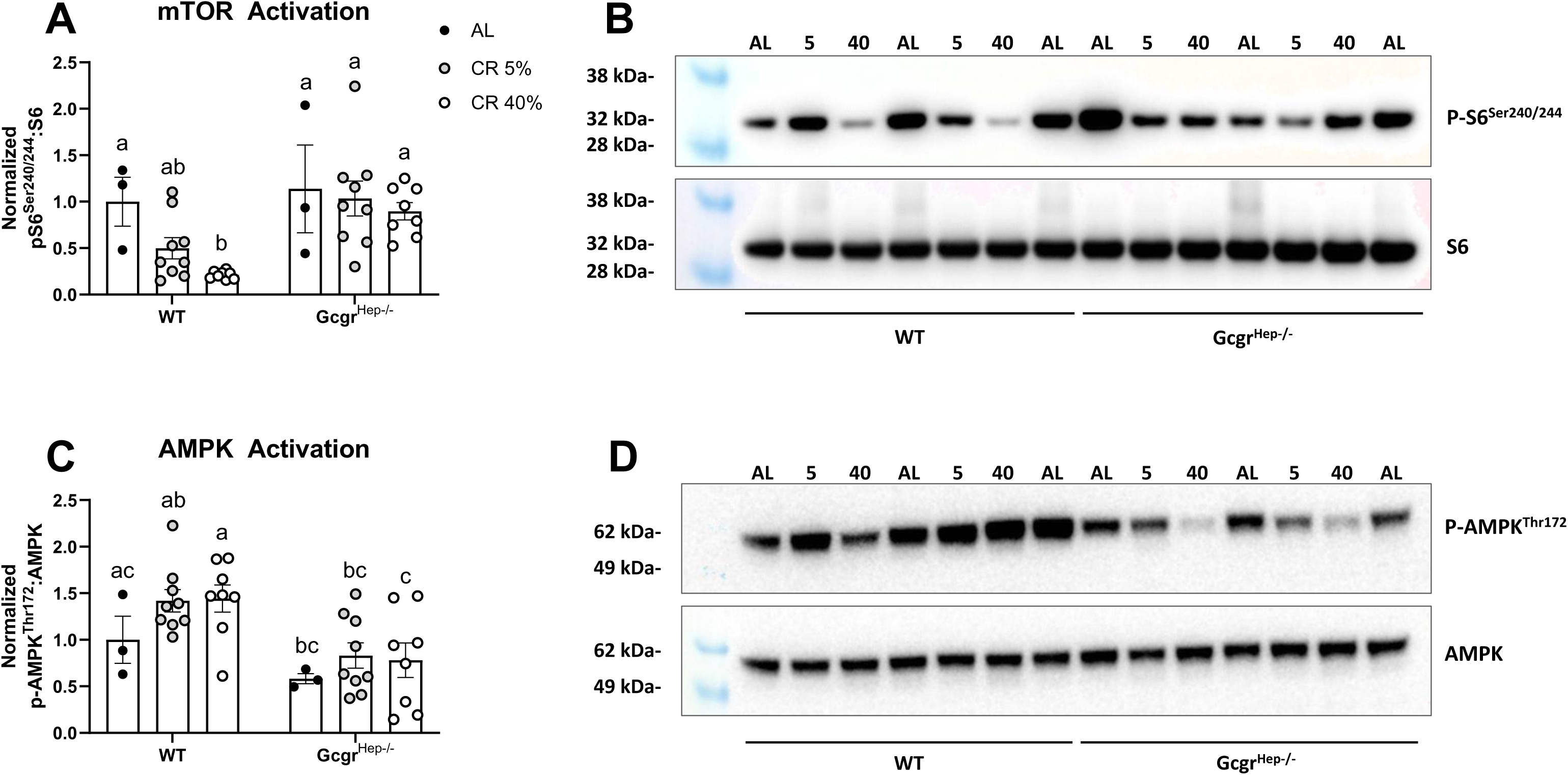
mTOR and AMPK activation at the liver. Western blot quantification of A-B) phosphorylated to total ribosomal S6 protein expression and representative blot, and C-D) phosphorylated to total AMPK protein expression at the liver in 17-month-old liver specific glucagon receptor knockout mice (Gcgr^hep-/-^) and wildtype control littermates (WT) fed an ad libitum (AL), 5% calorie restricted (CR 5%), or 40% calorie restricted (CR 40%) diet, initiated at 4 months of age) n= 3-9 mice per group. ^a,b^Superscript letters that differ indicate differences, P< 0.05; two-way ANOVA with Tukey’s adjustment for multiple comparisons, data are means ± SEM.

### 3.5 Pharmacologic Glucagon Receptor Agonism decreases mTOR activity at the liver

Having shown that Gcgr^Hep-/-^ mice fail to respond to CR with a decrease in mTOR activity at the liver, we next set out to determine if exogenous glucagon treatment could affect mTOR activity at the liver in both young adult (6 months of age) and aged (17 months of age) wildtype male C57BL/6J mice. After 4 weeks of treatment (three times per week) with a long-acting glucagon analogue, liver mTOR activity, as assessed by the ratio of phosphorylated (Ser^444/440^) to total ribosomal S6 protein, decreased in both young adult (P=0.005, **Figure 6A**) and aged (P=0.047, **Figure 6B**) mice compared to PBS treated control mice (Full blots provided in Supplemental Figure 9).

**Figure 6:**
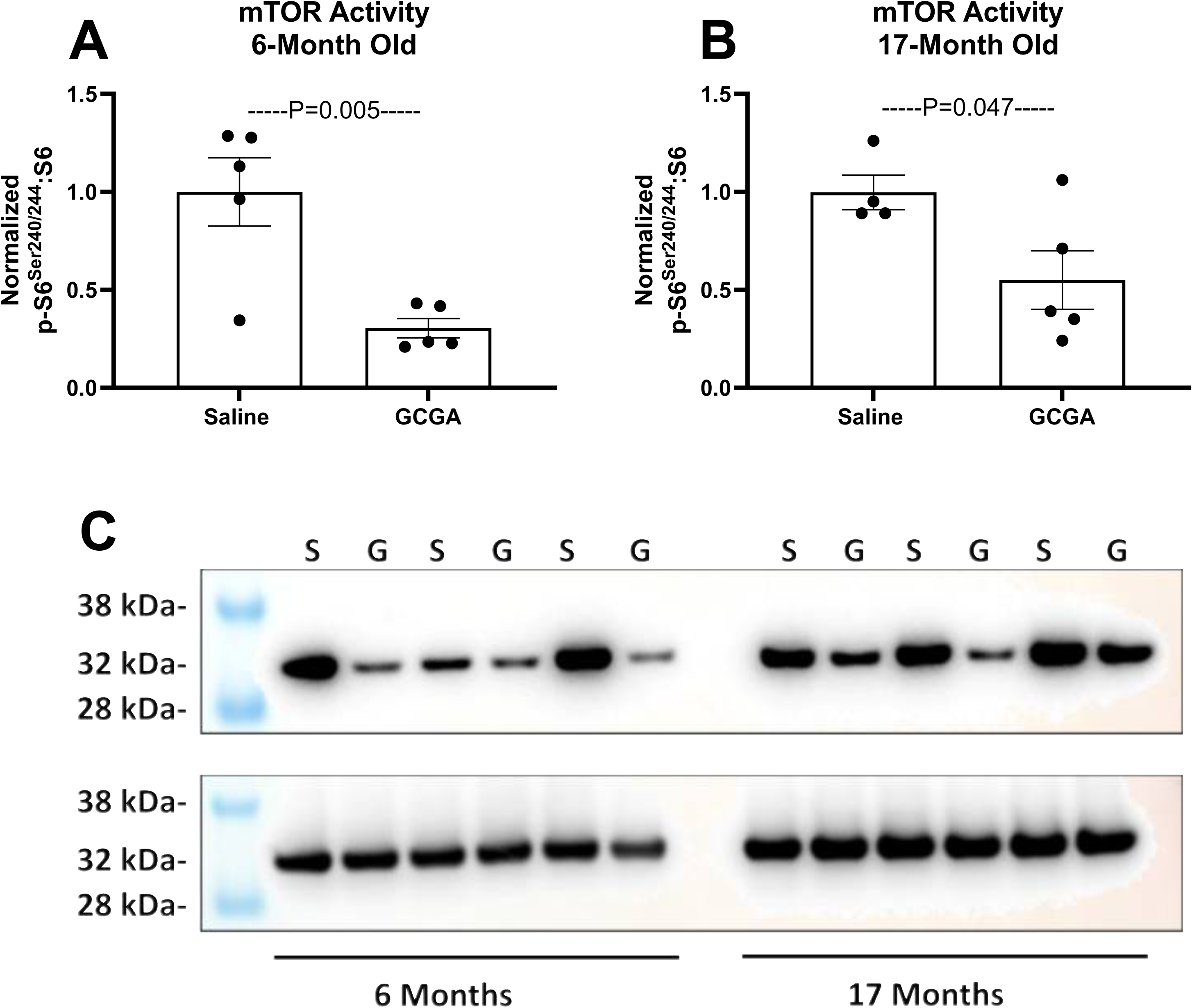
Treatment with a long-acting glucagon agonist inhibits mTOR activity. Western blot quantification of phosphorylated to total ribosomal S6 protein expression at the liver in wildtype male C57BL/6J mice treated with a long-acting glucagon analogue (GCGA, 3 nmol/kg BW subcutaneous, three times per week for 4 weeks). Tissues were collected from A) 6-month-old and B) 17-month-old mice. C) representative blot. S: PBS, G: GCGA, n= 5 mice per group; unpaired t-test, data are means ± SEM.

### 3.6 Caloric Restriction fails to improve Physical function in Gcgr KO mice

Chronic lifelong CR and prolonged fasting improves physical function in aging rodents [48, 49] [34]. In line with these findings, we found that 15% CR (initiated at 4 months of age) improves physical function (balance and coordination, as assessed by rotarod time to fall) in WT littermates mice at both 6 and 17 months of age. Yet we observed a diet*genotype interaction (6 month: P=0.03, 17 month: P=0.04), as CR fails to improve physical function in Gcgr KO mice (**Supplemental Figures 7A-B**).

## 4. Discussion and Conclusion

To understand the accelerated aging caused by obesity and the enhanced longevity resulting from CR, researchers have focused largely on the role of insulin and insulin signaling [50–54]. As such, they have established that genetic or dietary interventions that decrease circulating insulin enhance lifespan and decrease age-related diseases in the mouse [6, 55–57]. In the present study, we set out to understand the role of glucagon, the counterregulatory hormone to insulin, in normal lifespan and the healthspan benefits of calorie restriction in male mice. In so doing, we identified a critical role for glucagon receptor signaling in maintaining normal lifespan in chow-fed and high fat diet-fed mice and found that glucagon receptor signaling is indispensable for the healthspan benefits of lifelong 15% caloric restriction initiated at 4 months of age.

Lowering circulating insulin and enhancing insulin sensitivity extends lifespan in C. elegans [1, 2], drosophila [3, 4], and mice [6, 7]. Our lifespan studies demonstrate that, although global deletion of the glucagon receptor and high-fat diet feeding conditions. Further, we observed that calorie restriction does not extend survival at 19 months of age in Gcgr KO mice. This finding that glucagon receptor signaling is necessary for normal lifespan across dietary paradigms (low-fat chow fed, HFD induced obesity, and calorie restriction) suggests that glucagon receptor signaling regulates aging.

Our necropsy data from mice in these lifespan studies are limited to only chow-fed Gcgr KO mice, as we collected necropsy data from only a subset of chow-fed Gcgr KO mice Thus, we cannot make a conclusive observation as to the cause of lifespan curtailment in these animals. Our limited necropsy data indicates several incidences of islet cell carcinoma in Gcgr KO mice. Thus, it is highly possible the that the alpha cell hyperplasia commonly observed in both global and liver-specific glucagon receptor knockout mice [31, 32] may lead to islet cell carcinoma in these animals. However, few Gcgr KO deaths occurred prior to middle age. In fact, only one Gcgr KO mouse died prior to 12 months of age. Moreover, there was no difference in lifespan between Gcgr KO mice with or without islet cell carcinoma at time of death. Thus, it is unlikely that the development of islet cell carcinoma would explain our observation that Gcgr KO mice at 6 months of age fail to respond to CR with improvements in both metabolic and physical function, or that liver-specific deletion of Gcgr prevents CR induced suppression of mTOR signaling and lowers AMPK activity at the liver in aged 17-month-old mice. These findings, together with our observation that pharmacologic glucagon receptor activation decreases mTOR activity in both young and aged C57BL/6J wildtype mice, propose that the effects of glucagon receptor signaling in healthspan, aging, and the response to CR are not limited to islet cell carcinoma.

Beyond lifespan extension, caloric restriction without malnutrition enhances healthspan, decreasing age-related disease and improving metabolic and physical function in laboratory rodents and non-human primates [34, 58–63]. We demonstrate that 15% calorie restriction initiated at 4 months of age improves both metabolic and physical function at 6 and 17 months of age in WT littermates but fails to improve healthspan in Gcgr KO mice. Glucagon receptor deletion in the mouse was initially studied as a model for exploring the potential to lower glucose through glucagon receptor antagonism and prevent hyperglycemia associated with β-cell destruction [64–66]. We confirm these previous landmark studies demonstrating the glucose- and insulin-lowering effect of global glucagon receptor deletion and glucagon receptor antagonism in the mouse. Despite these improvements glucose or insulin (Figure 3). Moreover, Gcgr KO mice are resistant to CR induced improvements in lipid homeostasis at both 6 and 17 months of age (Figure 4).

Previous studies have demonstrated the critical role of liver glucagon receptor signaling in the maintenance of lipid homeostasis, particularly in response to fasting [8]. Longuet and colleagues demonstrated that global deletion of the glucagon receptor in mice increases hepatic triglyceride secretion and inhibits fatty acid oxidation at the liver [8]. More recently, studies have shown the potential of glucagon containing di- and tri-agonists to treat obesity by lowering liver fat and improving lipid homeostasis in obese mice [26–28] and humans [29, 30]. Similarly, our studies highlight a critical role for glucagon receptor signaling in the lipid lowering effects of chronic caloric restriction in the mouse (Figure 4) at both young adulthood (6 months of age) and advanced age (17 months of age).

We imposed a 15% restriction to study the impact global glucagon receptor signaling in CR induced improvements in healthspan, a level of restriction lower than most studies investigating the impact of CR on healthspan and aging in mice, which typically employ a 30-40% caloric restriction [62, 63, 67]. Our previous work showed that a 15% CR improved both metabolic and physical function in aging mice [34], demonstrating similar improvements in both lipid and glucose homeostasis as we observed in WT littermate control mice in the present study. Given that we provided feed once daily prior to the onset of the dark cycle and mice consumed all feed within 2-3 hours, all calorie restricted mice underwent an extended fast of approximately 21-22 hours before being provided feed again. The healthspan benefits of time restricted feeding and intermittent fasting versus caloric restricted has recently been extensively studied [49, 68, 69] and reviewed [70, 71], with mounting evidence pointing to the healthspan benefits of intermittent fasting in both mice and humans. Interestingly, Duregon and colleagues (2023) [49] observed that time restricted feeding in aging female C57BL/6J mice resulted in a 15% reduction in caloric consumption, identical to the level of restriction in our present study. Of note, the 15% CR implemented in this study is similar to the 11.9 ± 0.7% calorie restriction achieved in participants from the CALERIE trial (Comprehensive Assessment of Long-Term Effects of Reducing Intake of Energy), the first ever long term (2 year) CR intervention in non-obese humans. In this clinical trial, although participants did not undergo extended periods of fasting, 11.9 ± 0.7% CR reduced LDL cholesterol, total: HDL

We studied two levels of restriction when examining AMPK and mTOR activity in Gcgr^hep-/-^ and wildtype littermate mice (Figure 5). Including both 5% and 40% CR groups to compare to AL-fed mice allowed us to assess differences between prolonged fasting only (5% CR) and prolonged fasting in combination with a robust caloric restriction (40% CR). As in our experiments imposing a 15% CR, both 5% and 40% restricted mice consumed their feed within 2-3 hours and, thus, underwent an extended fast of approximately 21-22 hours before again being provided feed. Interestingly, we show that an extended fast alone (5% CR) is not sufficient to significantly decrease mTOR activity at the liver in WT littermates (Figure 5A-B). This is in agreement with the findings of Duregon and colleagues (2023) [49] and supports the hypothesis that increased fasting duration in combination with CR may promote greater geroprotection than extended fasting alone.

Whether chronic CR, as opposed to a single acute fast, activates AMPK at the liver is debated. While some studies show that CR activates AMPK in rat liver [74] and mouse heart [75] and skeletal muscle [76], others show no effect of chronic CR on liver, heart, or skeletal muscle AMPK activation in the mouse [77]. In fact, Kazuo and colleagues reported a decrease in liver AMPK activation in response to chronic 30% CR and no change in AMPK activation in response to alternate day fasting in Wistar rats [78]. In line with the latter, we found no effect of a 5% or 40% CR on liver AMPK activation. Importantly, regardless of diet, liver-specific deletion of the glucagon receptor decreases AMPK activation at the liver (Figure 5C-D).

The AMPK and mTOR pathways play a critical role in normal aging and the healthspan extension resulting from CR. Previous studies have demonstrated that AMPK activity is required for CR-induced improvements in metabolic function [79, 80]. A decrease in the mTOR signaling pathway promotes longevity and improves healthspan and may be a critical mechanism by which caloric restriction extends lifespan and improves healthspan [81]. AMPK activity at the liver is increased by glucagon [12]. In fact, glucagon suppresses hepatic mTOR signaling [82] by activating AMPK [12]. In line with these studies, we show that deletion of the glucagon receptor specifically at the hepatocyte decreases liver AMPK activity and prevents CR-induced suppression of mTOR activity at the liver (Figure 5). Finally, we show that 4 weeks of treatment with a long-acting glucagon receptor agonist suppresses mTOR activity at the liver in both young and aged C57BL/6J wildtype mice (Figure 6). Together, these findings demonstrate that not only does glucagon signaling at the liver suppress mTOR signaling, but that it is necessary in mediating the suppressive effects of caloric restriction on mTOR activity and is required for normal levels of AMPK activity at the liver.

Our observation that CR fails to decrease liver Nr1d1 mRNA expression in Gcgr KO mice at both 6 and 17 months of age is in line with these findings (Supplemental Figures L&P). The nuclear receptor REV-ERBα, encoded by the Nr1d1 gene, activates mTORC1 in hepatocytes [83]. Furthermore, Verlande and colleagues (2021) show that glucagon can decrease REV-ERBα protein levels in liver via cAMP driven PKA activation and subsequent destabilization of the REV-ERBα protein [84]. These findings of Verlande and colleagues complement our findings that glucagon receptor signaling at the liver is both sufficient and necessary to mediate the suppressive effects of caloric restriction on mTOR activity, providing a potential mechanism by which glucagon receptor signaling regulates normal lifespan and mediates CR induced improvements in healthspan and lifespan.

## Limitations and Conclusion

As previously discussed, our ability to interpret the findings of the lifespan studies are limited because our necropsy data is limited to only AL fed Gcgr KO mice. A larger scale lifespan study with more thorough post-mortem histopathological analysis is required to gain a full understanding of the impact of glucagon receptor signaling in lifespan and the cause of death in mice lacking glucagon receptor signaling.

Despite the limitations of our lifespan data, findings from our cross-sectional studies in calorie restricted mice provide robust evidence that glucagon receptor signaling is required for the healthspan benefits of CR and that glucagon receptor signaling at the liver is required for the CR-induced decrease in hepatic mTOR signaling, a primary driver of aging [17–19, 46, 85]. Strengthening these findings, our agonist studies show that pharmacologic glucagon receptor agonism regulates the same nutrient sensing pathways at the liver that are responsive to CR. Altogether, our findings provide the first line of evidence suggesting that glucagon receptor signaling is a regulator of aging and plays a critical role in mediating the healthspan benefits of CR in both young adult and aging mice. Furthermore, our findings demonstrating the suppressive effect of pharmacologic glucagon receptor agonism on liver mTOR activity and propose further investigation into the potential for glucagon containing dual and tri agonists to slow aging. To expound upon these findings, future studies must investigate the tissue-specific effects of glucagon receptor signaling in aging, the role of glucagon receptor signaling in nutrient sensing pathways that regulate aging, and comprehensive lifespan studies to understand the potential role of glucagon receptor signaling in the development of age-related disease and cancer progression.

## Authors’ contributions

J.H.S. conceived and designed the research. K.R.B., I.R.B., T.J.M., A.V., S.G., and T.F. performed *in vivo* mouse studies, wet lab experiments, and corresponding data analysis. Y.S., T.W.C and J.Z. performed *in vivo* animal studies. H.W. and F.A.D. performed echo MRI body composition and indirect respiration calorimetry analyses. D.G.B. performed necropsies and histopathological analyses. D.J.D. generated the glucagon receptor flox mouse and provided feedback. K.R.B., I.R.B., and J.H.S. interpreted results of experiments. J.H.S wrote the manuscript with contributions from all authors.

## Acknowledgements

We thank Dr. Maureen Charron, PhD for providing the Gcgr KO mouse.

## Data availability

The datasets used and/or analyzed during the current study are available from the corresponding author on reasonable request.

## Supplemental Data Figure Legends

**Figure S1:**
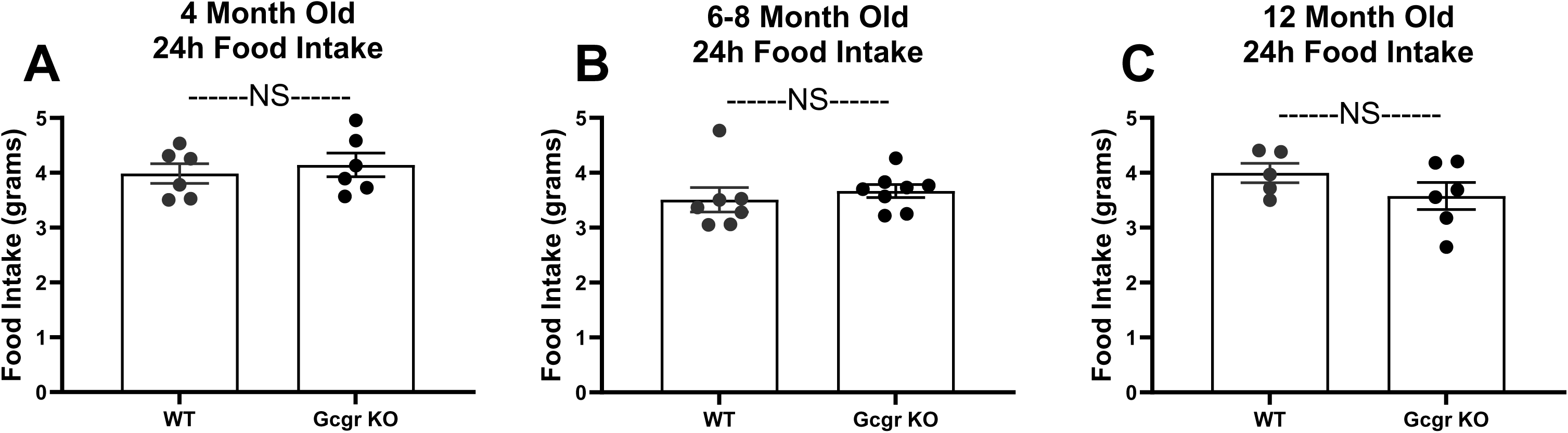
Food Intake. Ad libitum 24-hour food intake in WT and Gcgr KO mice at 4- (A), 6- (B), and 12- (C) months of age. Gcgr KO: global glucagon receptor knockout, WT: wildtype littermate controls. Independent t-test. NS, not significant, data are means ± SEM.

**Figure S2:**
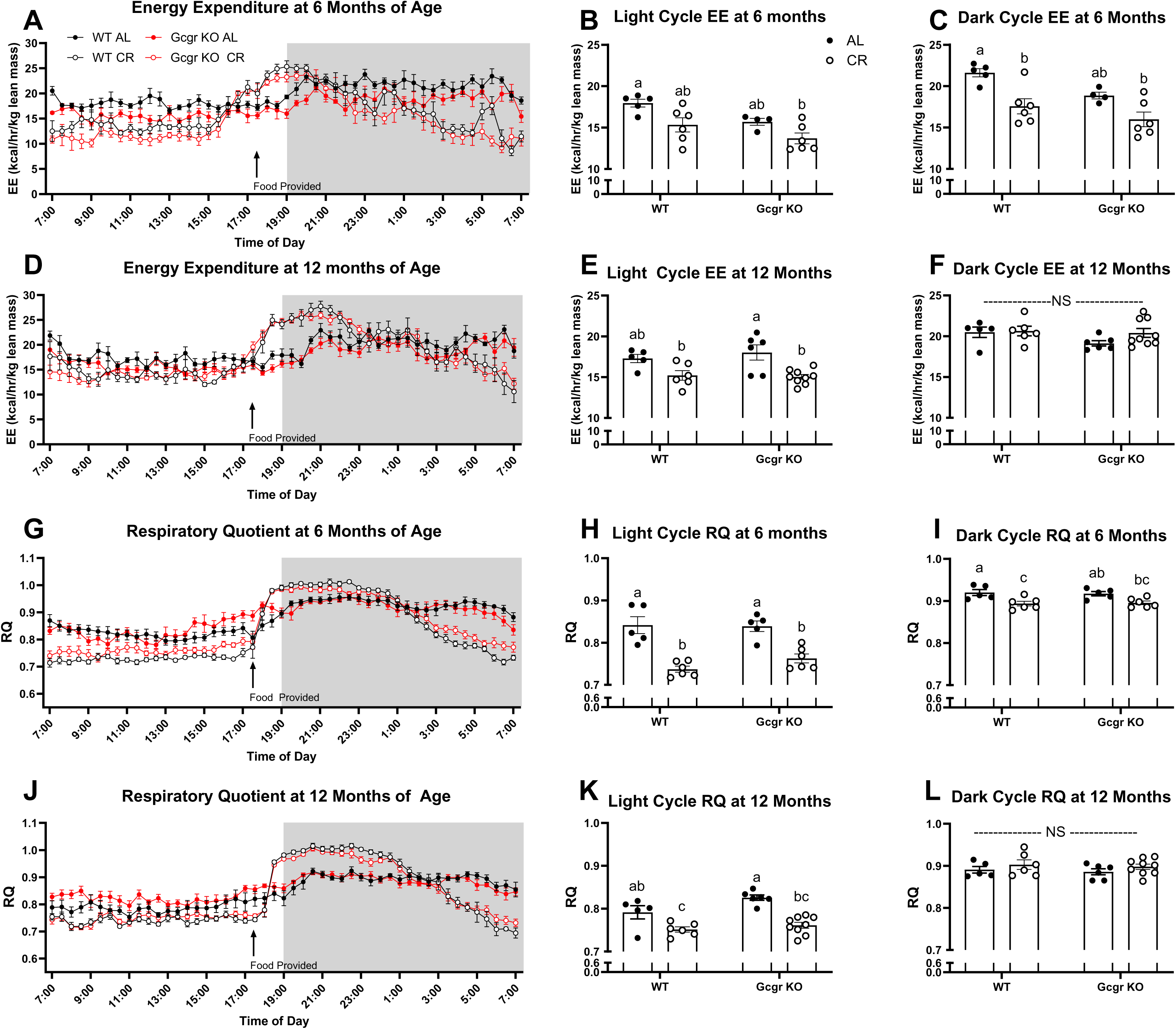
Whole Body Energy Expenditure and Substrate Utilization. Energy expenditure (EE, kcal/ hour/ kg lean mass) over 24 hours, light and dark cycle average energy expenditure in 6-month-old (A-C: n= 5-7 mice per group) and 12-month-old (D-F: n= 5-9 mice per group) mice. Corresponding respiratory quotient (RQ) in 6 month (G-I) and 12 month (J-L) old mice. Gcgr KO: global glucagon receptor knockout, WT: wildtype littermate controls. AL, ad libitum fed; CR, calorie restricted (15% initiated at 4.5 months of age). ^a,b^Superscript letters that differ indicate differences, P≤ 0.01; two-way ANOVA with Tukey’s adjustment for multiple comparisons, data are means ± SEM.

**Figure S3:**
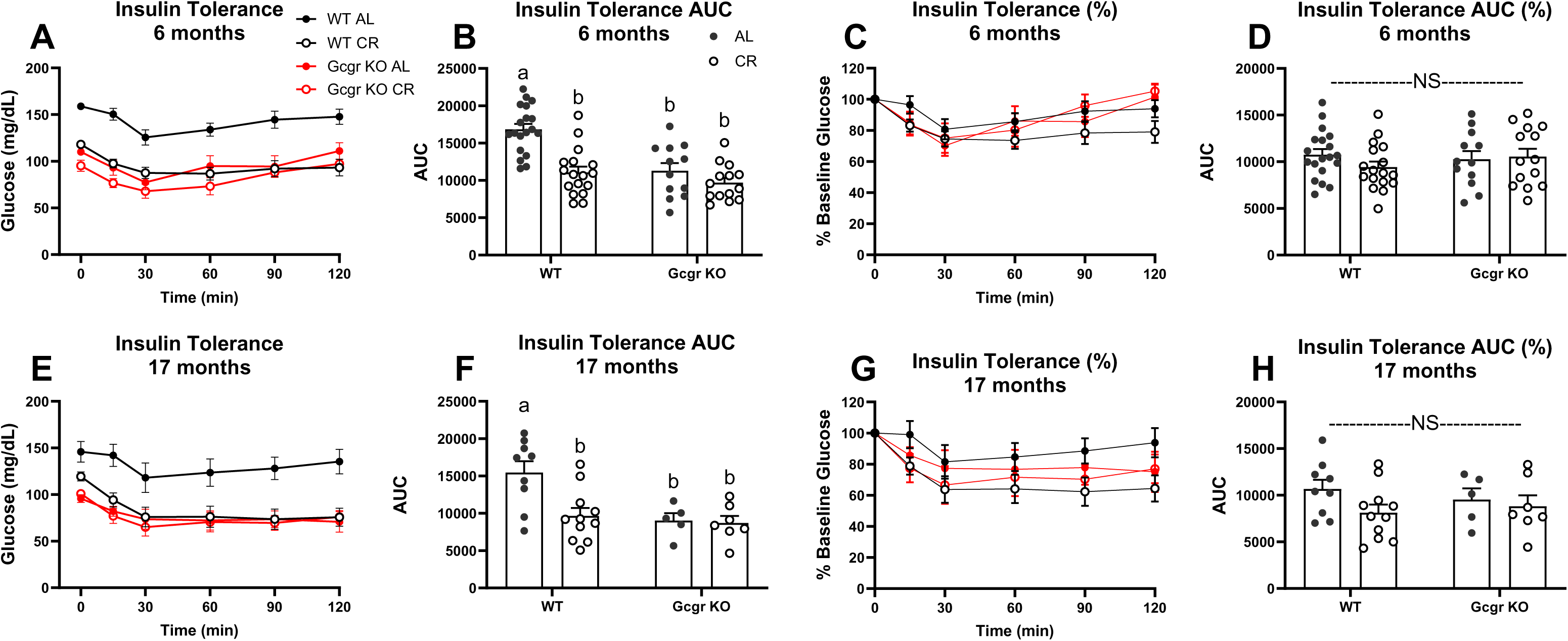
Insulin Tolerance. Insulin sensitivity, as assessed by an insulin tolerance test (0.5 IU/kg, intraperitoneal) at 6- (A-D) and 17- (E-F) months of age. Ad libitum (AL), 15% calorie restricted (15% CR). ^a,b^Superscript letters that differ indicate differences, P≤ 0.01; two-way ANOVA with Tukey’s adjustment for multiple comparisons. NS, not significant, data are means ± SEM.

**Figure S4:**
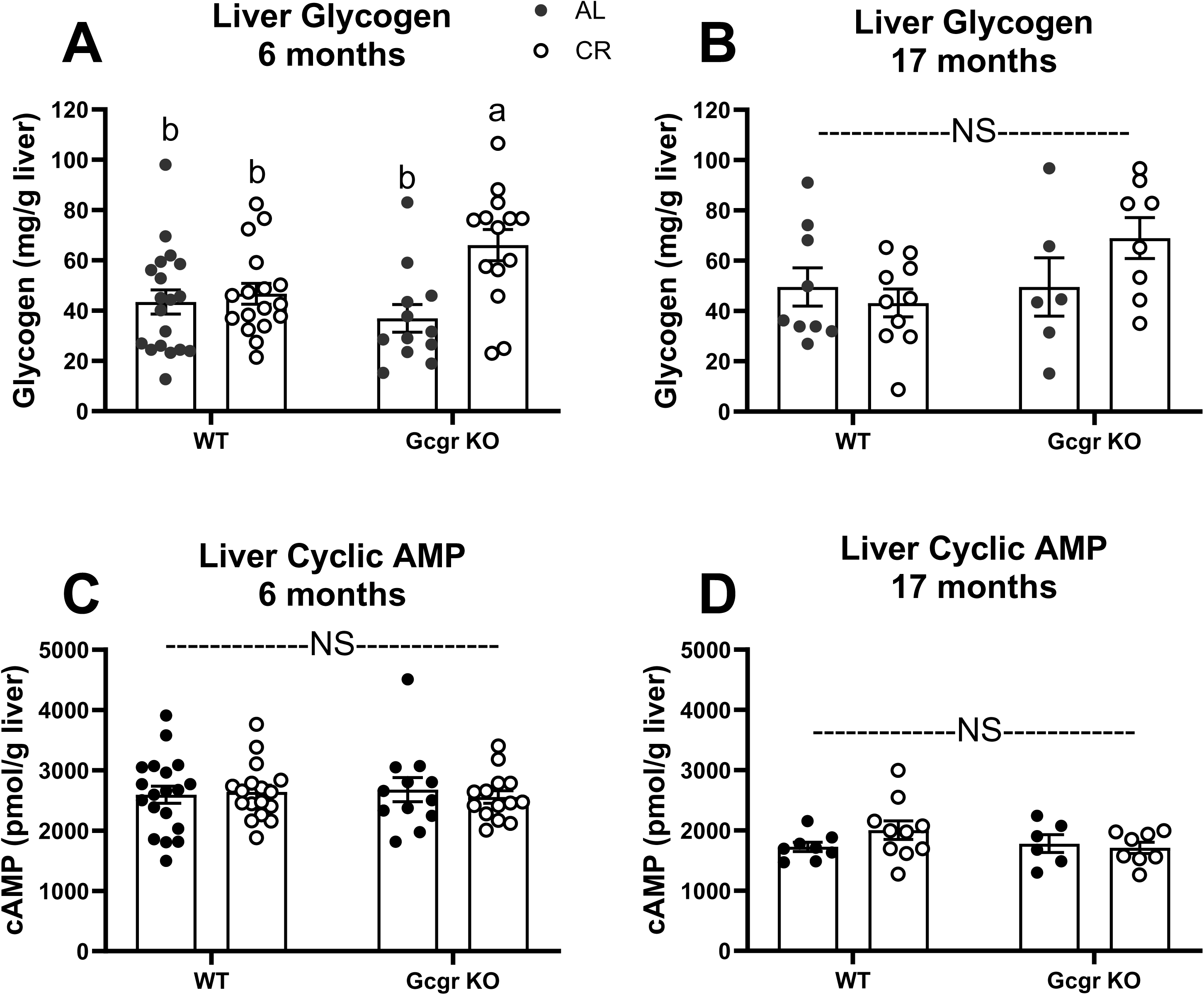
Liver Glycogen and cyclic AMP content. Liver glycogen (mg/g liver) in 6- (A) and 17- (B) month old mice and liver cyclic AMP (cAMP) in 6- (C) and 17- (D) month old mice. AL, ad libitum fed; CR, calorie restricted (15% initiated at 4.5 months of age). Gcgr KO: global glucagon receptor knockout, WT: wildtype littermate controls. ^a,b^Superscript letters that differ indicate differences, P< 0.05; two-way ANOVA with Tukey’s adjustment for multiple comparisons, data are means ± SEM.

**Figure S5:**
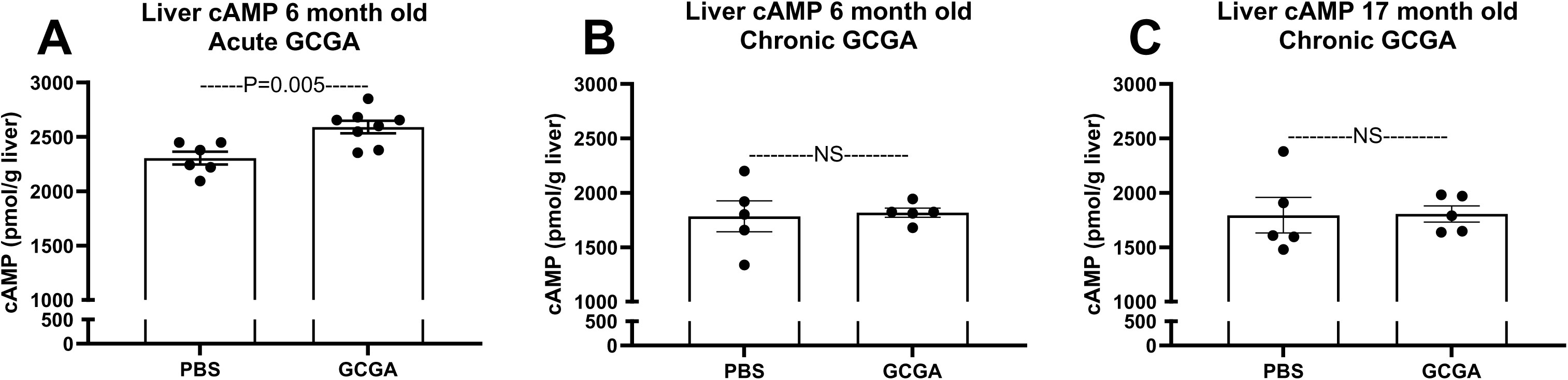
Glucagon receptor agonism only increases liver cAMP acutely. Liver cyclic AMP (cAMP) content in wildtype male C57BL/6J mice treated with a long-acting glucagon analogue A) Acutely (Single does GCGA, 1.5 nmol/kg BW subcutaneous) and B-C) Long-term (GCGA, 3 nmol/kg BW subcutaneous, three times per week for 4 weeks). Tissues were collected from 6-month-old (A-B) and 17-month old (C) mice. n= 5-7 mice per group; unpaired t-test, data are means ± SEM.

**Figure S6:**
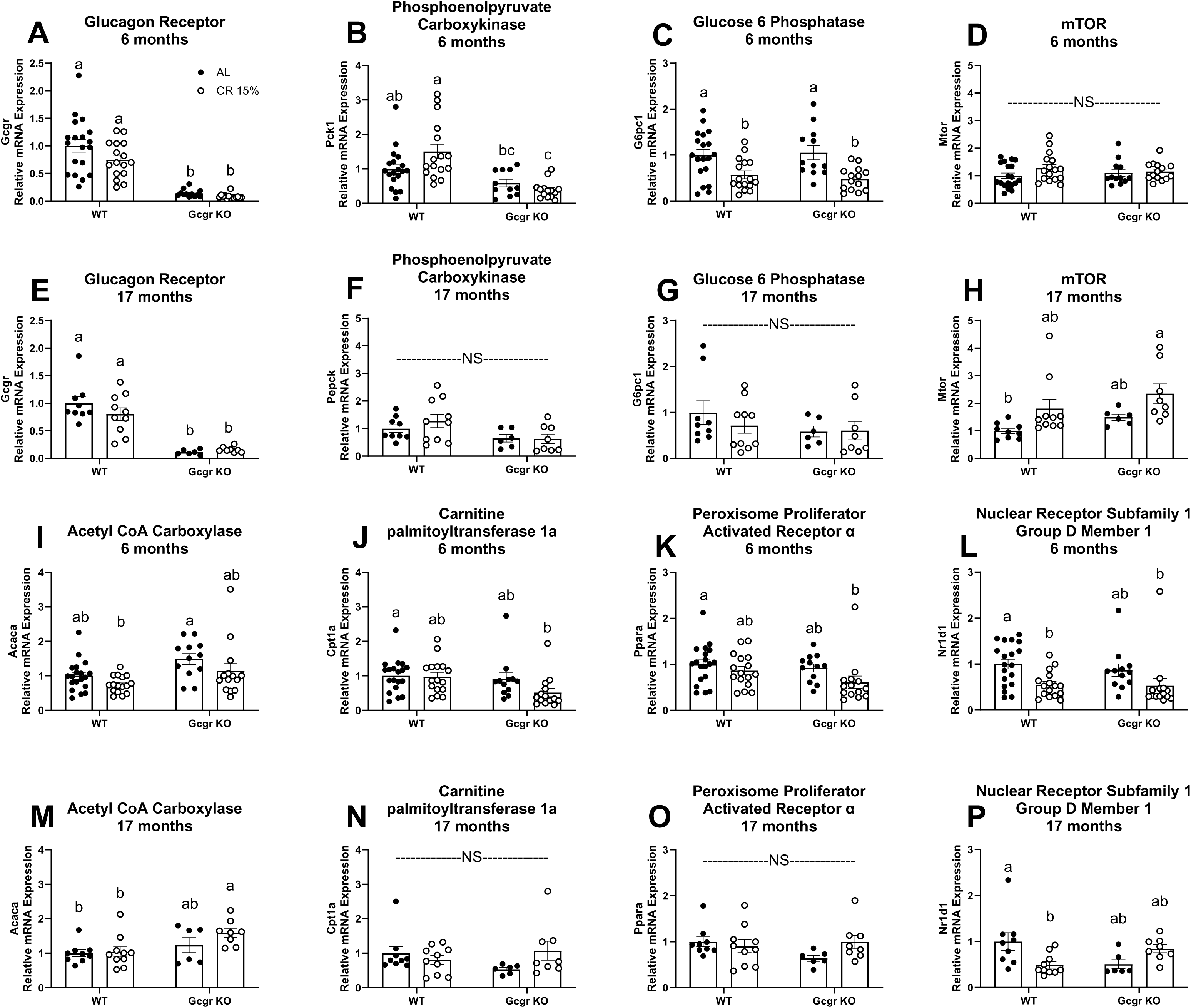
Hepatic Gene Expression. Liver mRNA expression of glucagon receptor (Gcgr, A&E), phosphoenolpyruvate carboxykinase (Pck1, B&F), glucose 6 phosphatase (Gcpc1, C&G), mechanistic target of rapamycin (Mtor, D&H), acetyl-Coenzyme A carboxylase alpha (Acaca, I&M), carnitine palmitoyltransferase 1a (Cpt1a, J&N), peroxisome proliferator activated receptor alpha (Ppara, K&O), and nuclear receptor subfamily 1, group D, member 1 (Nr1d1, L&P). ^a,b^Superscript letters that differ indicate differences, P≤ 0.01; two-way ANOVA with Tukey’s adjustment for multiple comparisons. NS, not significant, data are means ± SEM.

**Figure S7:**
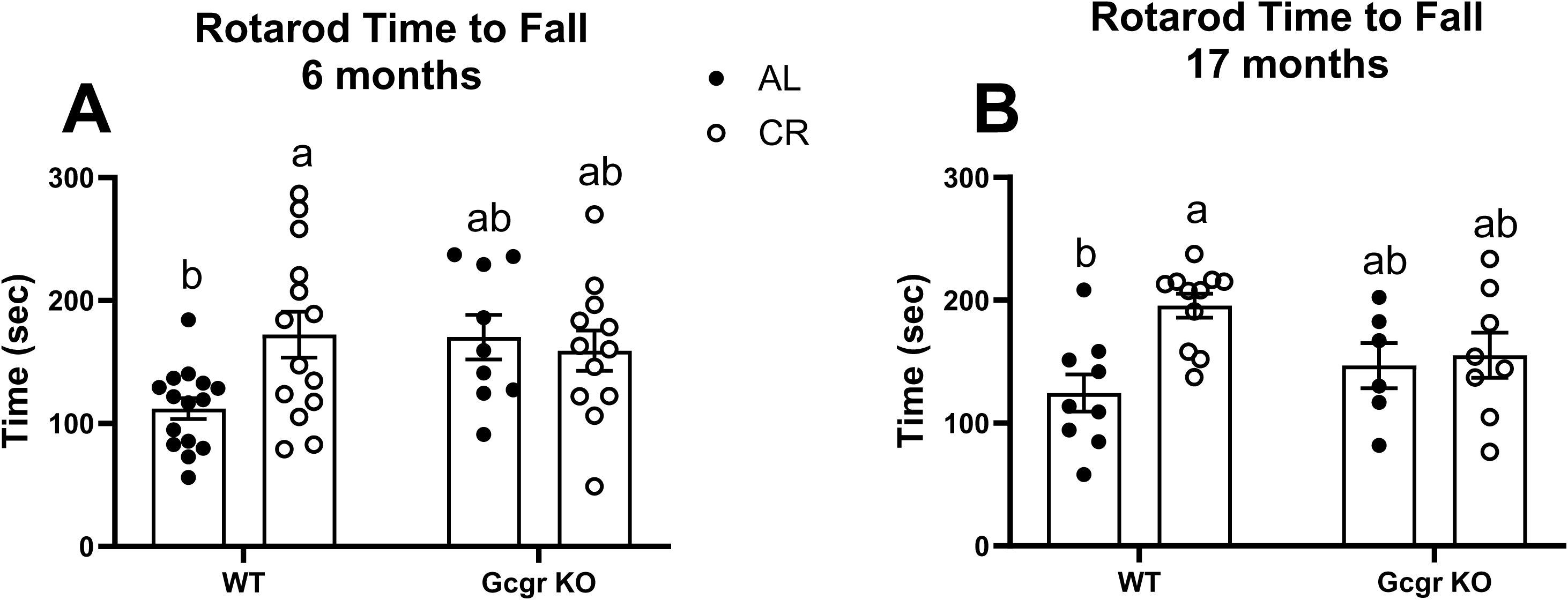
Physical Function. Physical function as assessed by time to fall in seconds on a Rotarod performance test in 6- (A) and 17- (B) month old mice. AL, ad libitum fed; CR, calorie restricted (15% initiated at 4.5 months of age). Gcgr KO: global glucagon receptor knockout, WT: wildtype littermate controls. ^a,b^Superscript letters that differ indicate differences, P< 0.05; two-way ANOVA with Tukey’s adjustment for

**Figure S8:**
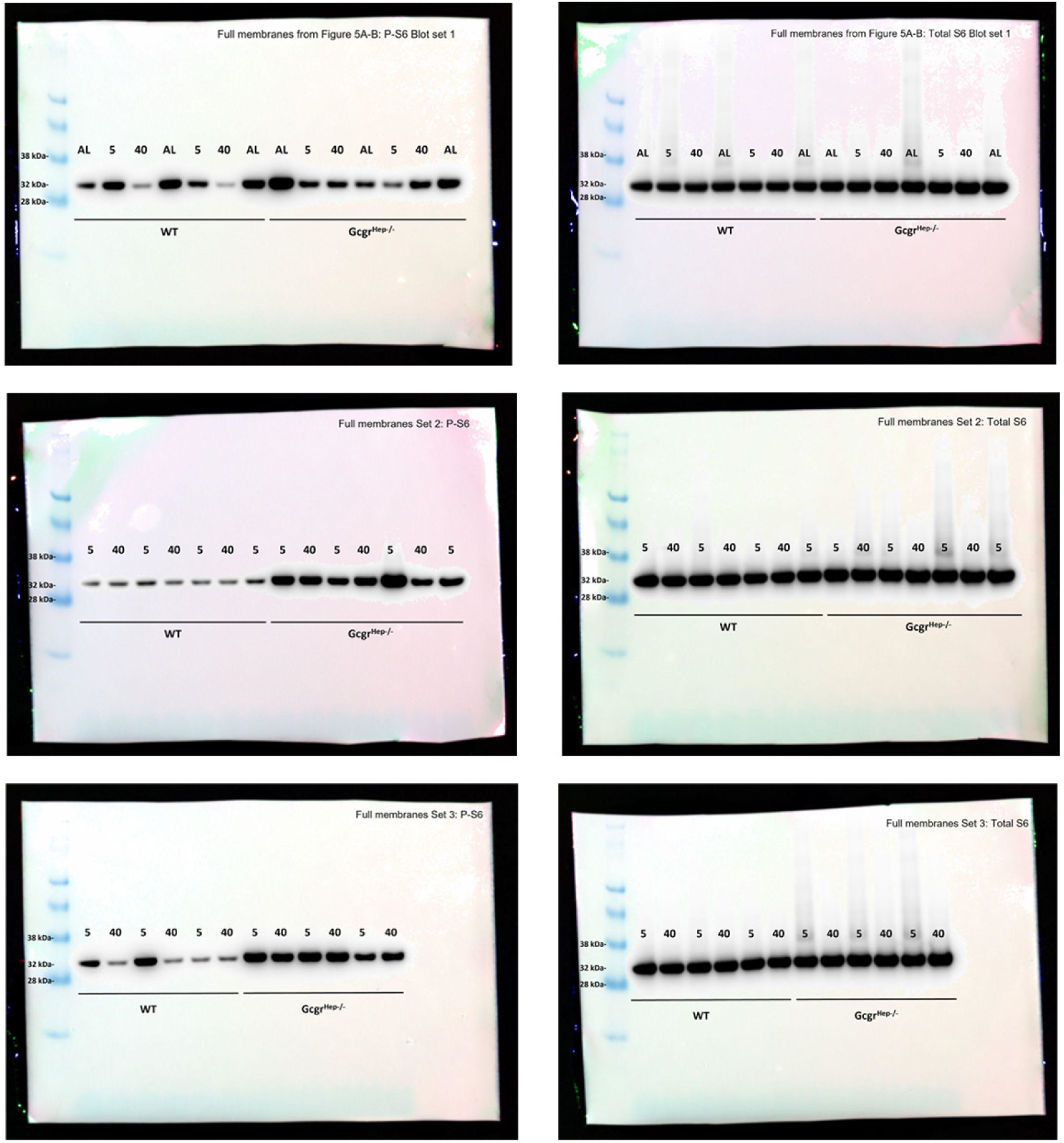
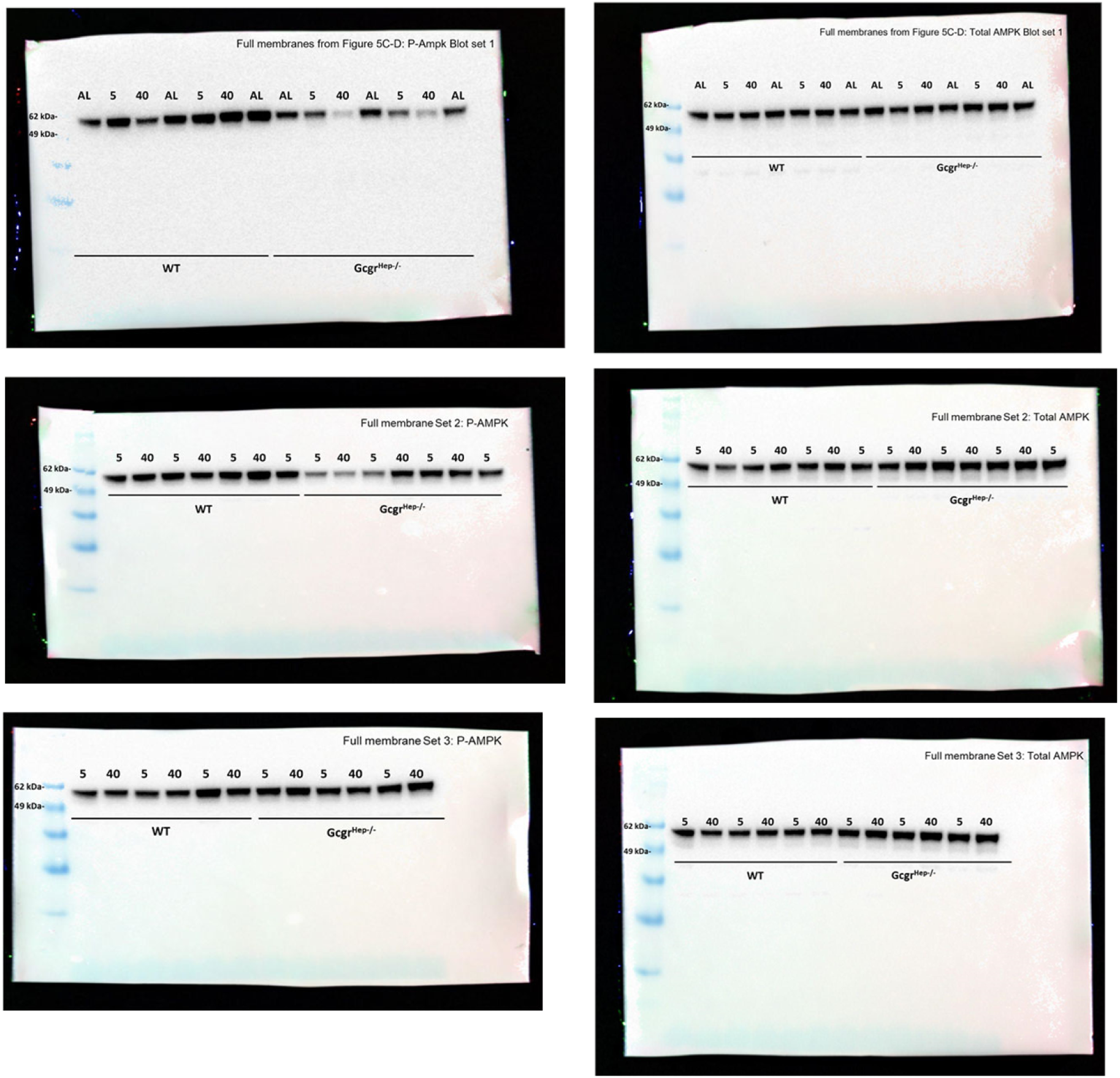
A) Full Western Blot Membranes from Figures 5C-D. Phosphorylated and total Ribosomal S6 Protein blots. B) Full Western Blot Membranes from Figures 5A-B. Phosphorylated and total AMPK blots.

**Figure S9:**
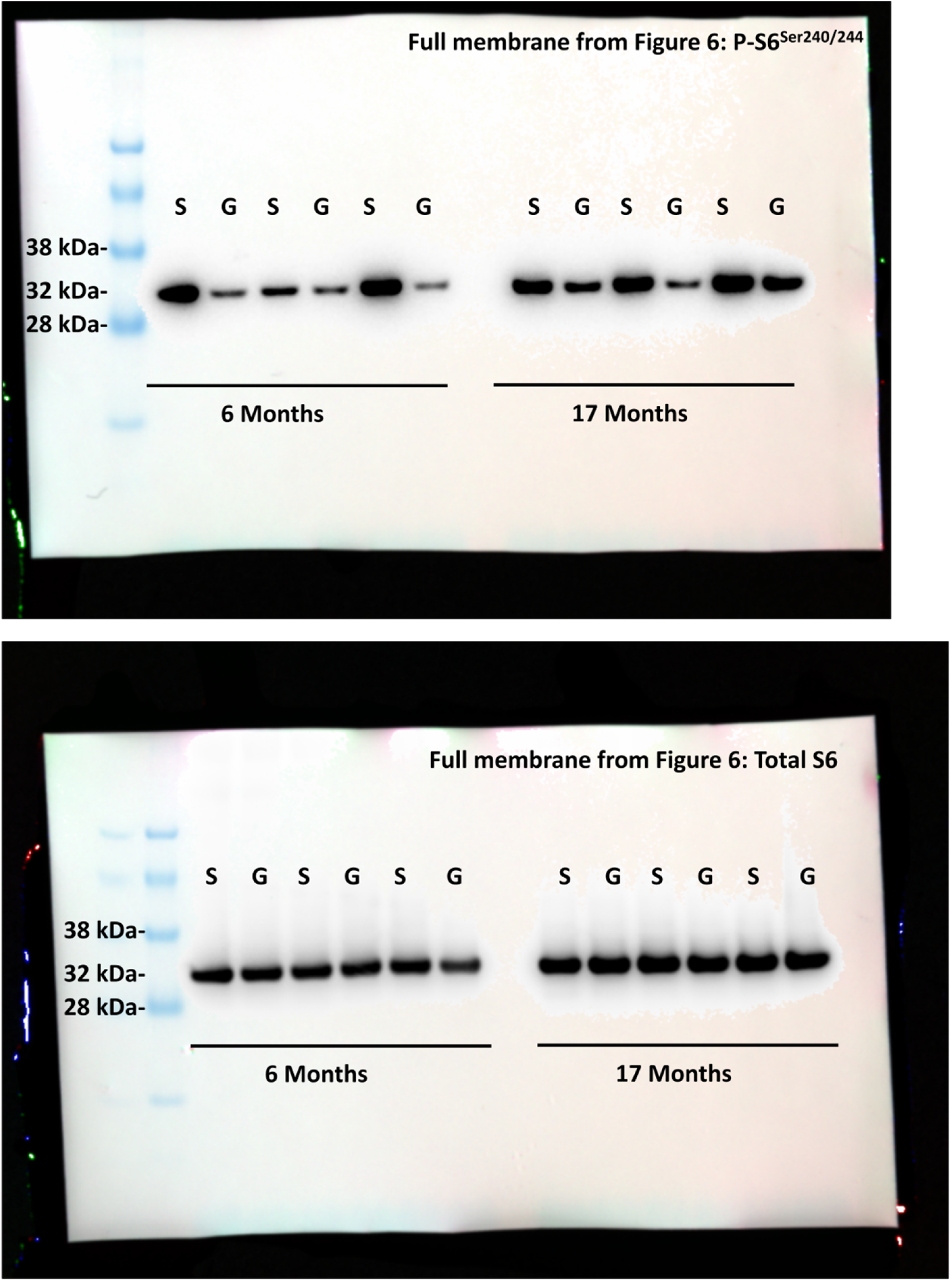
Full Western Blot Membranes from Figure 6, Phosphorylated and total Ribosomal S6 Protein blots.

**Supplemental Data Table 1.**

| Supplemental Data Table 1. Pathology at Time of Death in Gcgr KO mice | Gcgr KO (n=9) |
| --- | --- |
| **Pancreas** |  |
| Islet cell carcinoma | 5 |
| **Liver** |  |
| Hepatic Metastasis of Islet cell carcinoma | 1 |
| **Kidney** |  |
| Moderate glomerular amyloidosis | 1 |
| Mild to moderate post-mortem autolysis | 3 |
| **Heart** |  |
| None |  |
| **Lung** |  |
| Mild Focal alveolar histiocytosis | 1 |
| **Stomach** |  |
| Wall Discoloration | 1 |
| Distension | 2 |
| **Cecum** |  |
| Enlarged | 1 |
| **Gallbladder** |  |
| None |  |
| **Spleen** |  |
| Extramedullary hematopoiesis | 1 |
| **Prolapse (of any organ)** |  |
| Rectal | 1 |
| Penile | 1 |
| **Abdomen** |  |
| Cranioventral Round cell neoplasia | 1 |
| Distension | 1 |
| **No significant lesions detected (including the middle ear)** | 3 |

